# Elucidating small RNA pathways in *Arabidopsis thaliana* egg cells

**DOI:** 10.1101/525956

**Authors:** Stefanie Sprunck, Marc Urban, Nicholas Strieder, Maria Lindemeier, Andrea Bleckmann, Maurits Evers, Thomas Hackenberg, Christoph Möhle, Thomas Dresselhaus, Julia C. Engelmann

**Author notes:** authors to whom correspondence should be addressed, phone: +49/(0).

## Abstract

- Small RNA pathway components and small RNA profiles of flowering plant egg cells are largely unexplored, mainly because they are not easily accessible but deeply buried inside the ovary.
- We describe here the utilization of proliferating callus tissue that adopted transcriptome features of Arabidopsis egg cell as a tool to explore small RNA pathway components and small RNA profiles in egg cells. We furthermore complement our studies with mRNA-Seq data from isolated Arabidopsis egg cells and provide data validation by promoter-reporter studies and whole mount i*n situ* hybridization.
- Sequencing of small RNA libraries demonstrate the predominance of TE-derived siRNAs in the egg cell-related callus. TE-features and expression profiles suggest post-transcriptional silencing of activated *Gypsy*-like LTR retrotransposons, whereas the majority of class II DNA transposons belonging to *Copia, CACTA, hAT*-like and *Mutator* superfamilies are subjected to transcriptional silencing.
- Small RNA-seq furthermore led to the identification of differentially expressed known and novel miRNAs whose expression in the egg cell was verified by small RNA whole mount in situ hybridization. Both the strong expression of miRNAs in the egg-cell-adjoining synergids and the secretion of miRNAs into the micropyle suggest hitherto undescribed roles for these accessory cells in intercellular communication with the egg cell and the arriving pollen tube.
- In conclusion, our datasets provide valuable and comprehensive resources to study small RNA pathways and small-RNA-mediated epigenetic reprogramming during egg cell differentiation and the onset of plant embryogenesis.

## Introduction

The life cycle of flowering plants alternates between a dominant, diploid sporophytic phase and a short haploid gametophytic phase. Contrary to most animals, where the germline is set aside early during development and persists throughout the life history of the organism, flowering plant egg cells and sperm cells differentiate at a late stage of adult development, when flowers with reproductive organs are formed. Furthermore, flowering plant gametes are not direct products of meiosis but develop as part of a multicellular haploid gametophyte, where they are accompanied by accessory cells, some of which are pivotal in assuring successful double fertilization (Yadegari and Drews 2004; Higashiyama and Yang, 2017).

In most flowering plants, including Arabidopsis, female gametophyte development begins early in ovule development, when a sub-epidermal cell differentiates into the diploid megaspore mother cell which will initiate meiosis (megasporogenesis). After meiosis, only one megaspore develops into the female gametophyte (megagametogenesis). Whereas the other three megaspores undergo programmed cell death, this functional megaspore performs three free nuclear division cycles resulting in an eight-nucleate syncytium (Sprunck and Groß-Hardt 2011). Cell fate decisions are affected by positional cues, acting on the daughter nuclei in the syncytium according to their position along the distal (micropylar) to proximal (chalazal) axis (Skinner and Sundaresan, 2018). During the following cellularization, four cell types are established: The egg cell and two accessory synergid cells locate close to the micropylar entrance point of the pollen tube, two polar nuclei fuse and form the nucleus of the large central cell, whereas three antipodal cells locate to the chalazal pole of the female gametophyte. Thus, three major cell specification events occur during female gametophyte development: (*i*) the specification of the megaspore mother cell from a sporophytic precursor (somatic-to-reproductive cell fate transition); (*ii*) the selection of one haploid megaspore that will proceed with megagametogenesis; (*iii*) the specification of four distinct cell types during cellularization of the female gametophyte.

Over the past years, small RNA-mediated gene silencing has emerged as one of the most important mechanisms in regulating gene expression and repressing transposable elements (TEs) at the transcriptional or posttranscriptional level. Regulatory small non-coding RNAs include miRNAs (microRNAs), siRNAs (small interfering RNAs), animal-specific PIWI-interacting RNAs (piRNAs), as well as plant-specific tasiRNAs (*trans*-acting siRNAs) and phasiRNAs (phased secondary siRNAs). Core components of all small RNA-mediated silencing processes are Argonaute proteins (AGOs). Small RNAs are loaded onto AGOs to form multimeric effector complexes, which mediate sequence-specific post-transcriptional gene silencing the RNA level, or transcriptional gene silencing at the chromatin level (Borges and Martienssen, 2015; Feng and Qi, 2016; Nonomura, 2018).

In the germline of mammals and insects, PIWI-clade AGOs and adaptive piRNA genomic clusters play important roles in repressing TEs but also in the regulation of endogenous gene expression programs regulating germline stem cell maintenance, gametogenesis, maternal-to-zygotic transition and embryonic patterning (Vagin et al., 2006; Brennecke et al., 2007; Rouget et al., 2010; Rojas-Ríos and Simonelig, 2018). Although flowering plant genomes lack the PIWI-clade AGOs and piRNAs, there is substantial evidence that members of the AGO clade and mobile small noncoding RNAs control the somatic-to-reproductive cell fate transition and meiotic progression during megasporogenesis (Nonomura et al., 2007; Olmedo-Monfil et al., 2010; Singh et al., 2011; Liu et al., 2016). Furthermore, somatic small RNA pathways also promote gametogenesis (Tucker et al., 2012) and chromatin remodeling during gametogenesis is accompanied by the activation of transposable elements (TEs) and by histone replacement and histone modifications, respectively (Ingouff et al., 2010; She and Baroux, 2015). However, the impact of sncRNAs on chromatin changes, the transcriptional landscape and egg cell identity remains to be explored.

Here, we describe the utilization of a callus-like Arabidopsis cell line with egg-related transcriptomic features as a tool to explore small RNA profiles and small RNA pathway components in egg cells. Egg cell-related cell lines were generated by expressing RKD2, a family member of the plant-specific RWP-RK DOMAIN CONTAINING (RKD) transcription factors, ectopically in Arabidopsis seedlings. After validating the previously described egg-like transcriptomic features (Köszegi et al., 2011), we performed Illumina sequencing to investigate the small RNA profile and transcriptome in comparison to that from hormone-induced control calli. We furthermore complemented our studies with transcriptome data from isolated Arabidopsis egg cells and validated expression patterns in the Arabidopsis egg cell by promoter-reporter studies and whole mount in situ hybridization. Our studies revealed differentially expressed TE-derived siRNAs in the egg cell-related callus, indicating selective silencing and reactivation of TEs, reminiscent to the male germline. Beside identifying known and novel miRNAs in the egg cell, our results suggest a yet unknown role for the accessory synergid cells in intercellular communication with the egg cell and the arriving pollen tube. Our comprehensive small RNA and transcriptome data sets provide valuable resources to investigate small RNA pathways during egg cell differentiation and the onset of plant embryogenesis.

## Methods

### Plant materials and growth conditions

*Arabidopsis thaliana* ecotype Columbia (Col-0) was grown on soil under a short photoperiod (9 h of light, 21°C, 65% humidity) for 3 to 4 weeks, followed by a long photoperiod (16 h of light, 8,500 lux, 21°C, 65% humidity). To generate hormone-induced CIM callus, root explants were harvested from 2-week-old *Arabidopsis* seedlings (Col-0) grown in sterile culture on half strength MS medium supplemented with 2% (w/v) sucrose under a long photoperiod. 5 to 10 mm long root segments were cultured on callus-inducing medium (CIM) composed of 1x Gamborg’s B-5 medium B5 with 0.5 g/L MES, 20 g/L glucose, 1x Gamborg’s vitamin solution and 1% Phytoagar, supplemented with 2,4-D (final concentration 500 µg/L) and kinetin (final concentration 50 µg/L). Calli appeared after 7 to 10 days and were propagated in two-week subculture intervals on CIM medium.

### Cloning and generation of RKD2-induced callus

Pistils from flowers at developmental stage 12 (Smyth et al., 1990) were harvested to purify mRNA and generate cDNA as described previously (Sprunck et al., 2012). To generate callus lines with egg cell-like expression profile, the coding sequence of RKD2 (AT1G74480) was amplified from pistil cDNA using the primer pair RKD2fw /RKD2rev (Table S1), cloned into pENTR/D-TOPO (Thermo Fisher Scientific) and subsequently transferred into the GATEWAY-compatible destination vector pH7FWG2.0 (Karimi et al., 2002) via LR clonase reaction. The resulting expression vector *pH_35Sp:RKD2-GFP* was used for floral dip transformation of *Arabidopsis* (Clough and Bent, 1998). T1 seeds of transformed plants were surface-sterilized with 1% NaOCl and 70 % ethanol for 5 min each, washed 5 times with sterilized water and then plated on half-strength Murashige and Skoog (MS) medium with 2% (w/v) sucrose, 1% Phytoagar, and Hygromycin (30 µg/mL). RKD2-induced proliferating cells appeared 20 to 30 days later and were propagated in two-week subculture intervals on 1/2 MS, 2% (w/v) sucrose, 1% Phytoagar, Hygromycin (30 µg/mL). For RNA and protein work, callus material was collected with a sterile scalpel blade and immediately frozen in liquid nitrogen.

### Egg cell and central cell isolation

Arabidopsis egg and central cells were isolated from mature ovules according to a previously published protocol (Englhart et al., 2017). After isolation, cells were washed two times in mannitol solution as described, transferred to 500 µL Eppendorf^®^ LoBind tubes and immediately frozen in liquid nitrogen.

### Protein work

100 mg frozen callus material was ground to fine power, vortexed in 200 µL ice-cold extraction buffer (20 mM Tris pH 7.5, 300 mM NaCl, 5 mM MgCl2, 0.1% Triton X-100, 1x complete Proteinase-Inhibitor Cocktail from Roche, 50 µM MG-132) and incubated for ten minutes on ice. After centrifugation (10 minutes, 16.000 x g) the protein concentration of the clear supernatant was determined by the Bradford protein assay (Bio-Rad). After SDS-PAGE (7.5%), proteins were blotted on nitrocellulose (Amersham Protran 0.45 NC), stained with Ponceau S solution and blocked for one hour at room temperature in TBS-T (50 mM Tris/HCl pH 7.5, 150 mM NaCl, 0.2% Tween-20) supplemented with 5% milk powder. After washing in TBS-T (3x for 10 min) the membrane was incubated overnight at 4°C with the primary antibody. Polyclonal peptide antibodies against AGO proteins (Agrisera) were diluted as recommended by the supplier. Horseradish peroxidase (HRP)-coupled anti-rabbit IgG was used as secondary antibody. Luminol-based HRP-substrate (HRP juice, PJK GmbH) was used for detection.

### Total RNA and small RNA isolation for RNAseq

Total RNA (including low molecular weight RNA) was extracted from 200 to 300 mg of frozen callus, using the RNeasy Plant Mini Kit (Qiagen) and according to the manufacturer’s instructions. Three batches of each callus type (RKD2-induced callus and hormone-induced CIM callus, respectively) were used for RNA extraction and library preparation. For the RKD2-induced callus, sample 1 was generated from cell line 1 and samples 2 and 3 were generated from cell line 2. Purity and integrity of total RNA was assessed on the Agilent 2100 Bioanalyzer with the RNA 6000 Pico LabChip reagent set (Agilent). RNA extraction from egg cells was carried out at the genomics core facility of the University of Regensburg, Germany (Center for Fluorescent Bioanalytics (KFB); www.kfb-regensburg.de). For isolated egg cells, three replicates of 25 to 30 pooled egg cells were used for total RNA extraction according to the “Purification of total RNA from animal and human cells” protocol of the RNeasy Plus Micro Kit (Qiagen). In brief, RLT Plus buffer containing ß-mercaptoethanol was added to the frozen cells and the sample was homogenized by vortexing for 30 sec. Genomic DNA contamination was removed using gDNA Eliminator spin columns. Ethanol was added and the samples were applied to RNeasy MinElute spin columns followed by several washing steps. Total RNA was eluted in 12 µl of nuclease-free water.

### Library preparation and Illumina Next-Generation-Sequencing

Library preparation and deep sequencing was carried out by the KFB (Regensburg, Germany). For callus tissues, the same batch of total RNA was used for mRNA and small RNA library preparation, respectively. Poly(A)+ mRNA was selectively enriched from 500 ng total RNA using oligo-dT beads (NEBNext Poly(A) mRNA Magnetic Isolation Module; New England Biolabs, Ipswich, USA). The NEBNext Ultra RNA Library Prep Kit for Illumina (New England Biolabs) was used to prepare mRNA libraries, following the manufacturers guidelines. Small RNA library preparation was carried out according to the TruSeq Small RNA sample preparation guide (Illumina, Inc.), with the following modifications: For each of the six samples (three biological replicates), 1 µg RNA was used for ligation of the RNA 3’ and the RNA 5’ adapters, followed by reverse transcription to create single stranded cDNA. The cDNA was then PCR amplified for 12 cycles using a universal and an index primer. Following magnetic bead clean up (Agencourt AMPure XP, Beckman Coulter), small RNA species were gel-purified using an automated nucleic acid fractionation/extraction system with a collection width of 30 bp (LabChip XT DNA 300 Kit, Caliper Life Sciences). For egg cell libraries, the SMARTer Ultra Low Input RNA Kit for Sequencing v4 (Clontech Laboratories, Inc.) was used to generate first strand cDNA from 50 % of the extracted RNA. Double stranded cDNA was amplified by LD PCR (18 cycles) and purified via magnetic bead clean-up. Library preparation was carried out as described in the Illumina Nextera XT Sample Preparation Guide (Illumina, Inc.). 150 pg of input cDNA were tagmented by the Nextera XT transposome. The products were purified and amplified via a limited-cycle PCR program to generate multiplexed sequencing libraries. For the PCR step 1:5 dilutions of index 1 (i7) and index 2 (i5) primers were used.

### Illumina deep sequencing

Sequencing libraries were individually quantified using the KAPA SYBR FAST ABI Prism Library Quantification Kit (Kapa Biosystems, Inc.). Equimolar amounts of all libraries were pooled, and the pools were used for cluster generation on the cBot (TruSeq SR Cluster Kit v3). All sequencing runs were performed on a HiSeq 1000 instrument using TruSeq SBS v3 Reagents according to the Illumina HiSeq 1000 System User Guide. Callus and egg cell libraries were sequenced for 100 cycles (2×100nt, paired end) on two lanes. The six small RNA libraries were sequenced for 50 cycles (single end) on a single lane.

### Bioinformatic Analysis

FASTQC (http://www.bioinformatics.babraham.ac.uk/projects/fastqc/) was used to assess the quality of the different RNA-Seq libraries. Subsequently, low quality bases and adapter remnants were trimmed with Trimmomatic (Bolger et al. 2014) (small RNA parameters were ILLUMINACLIP:2:30:12 TRAILING:20 LEADING:20 MINLEN:18, searching for TruSeq Small RNA adapter remnants; mRNA parameters were ILLUMINACLIP:2:30:10:2:true TRAILING:3 LEADING:3 MINLEN:25, searching for Illumina universal paired-end and indexed adapters in the calli data, and for Nextera and SMARTer sequence remnants in the egg cell reads). The resulting reads were aligned to the *Arabidopsis thaliana* reference genome TAIR10 (http://www.arabidopsis.org/). Small RNA reads were mapped with butter (Axtell, 2014) with default parameters. Butter distributes reads mapping to multiple loci in the genome by an iterative process relying on the density distribution of uniquely aligned reads, and both unique and multi-mapping reads were considered. Alignment of mRNA reads was performed with Tophat2.0 (Kim et al., 2013) indicating an unstranded sequencing protocol and fragment length parameters retrieved from Bioanalyzer profiles (for callus mRNA: --mate-inner-dist 40 --mate-std-dev 40, for egg cell pool mRNA: --mate-inner-dist 190 --mate-std-dev 100). The annotation was retrieved from EnsemblPlants (ftp.ensemblgenomes.org, Arabidopsis_thaliana. TAIR10.22.gtf) (Kersey et al, 2014).

On the small RNA dataset, count tables required for differential expression analysis were produced using R/Bioconductor [GenomicAlignments] (Anders et al., 2015) considering only reads of lengths 18-24 nt. Read counts of transposable elements were retrieved from reads aligned to the antisense strand, while for all other features, reads mapping to the sense strand were counted. For quantifying mRNAs, featureCounts (Liao et al, 2014) was used counting uniquely mapped reads on both strands. For library size normalization and differential expression analysis the DESeq2 package (Love et al., 2014) was used independently for the small RNA and mRNA dataset. Genes with a false-discovery rate (termed p-value in the text) smaller than 0.00001 were regarded differentially expressed in the mRNA dataset comparing RKD2 to CIM calli. For Gene Ontology term analysis R/Bioconductor package ’goseq’ was used with annotations from GoMapMan (Ramsak et al., 2014). Transcripts per million (TPMs) were calculated for all mRNA samples for gene expression level comparisons. To visualize them in heatmaps, we used logarithmized transcripts per million (TPM) plus a pseudocount (log2[TPM+0.5]), as these were approximately Gaussian distributed, and normalized for between-sample variation. Prediction of novel miRNAs was performed with miR-PREFeR, and predicted mature miRNAs were included in the annotation file (Lei & Sun, 2014).

### Prediction of small RNA targets

Target predictions for miRNAs identified in our small RNA libraries were performed with psRNAtarget (http://plantgrn.noble.org/v1_psRNATarget/) (Dai & Zhao, 2011). Information on validated targets for miRNAs was retrieved from a comprehensive list provided by Konika Chawla (ftp://ftp.arabidopsis.org/home/tair/Genes/SmallRNAsCarrington/) and from miRTarBase 7.0 (http://mirtarbase.mbc.nctu.edu.tw/php/download.php).

### (Quantitative) RT-PCR

For RT-PCR, fifteen isolated cells (egg cells and central cells, respectively) were pooled for mRNA isolation and cDNA synthesis as described previously (Sprunck et al., 2005). For quantitative PCR, total RNA was extracted from RKD2-induced callus and hormone-induced CIM callus as described above and treated with DNAse I (Thermo Fisher Scientific). Poly(A)+ RNA was isolated from 5µg total RNA using the Dynabeads^®^ mRNA DIRECT™ Purification Kit (Thermo Fisher Scientific) with Dynabeads Oligo(dT)_25_. SuperScript III Reverse Transcriptase (Thermo Fisher Scientific) was used for first-strand cDNA synthesis following the manufacturers’ protocols with the addition of 1 µl Oligo(dT)_18_ primer and 1 µl RiboLock™ Ribonuclease Inhibitor (MBI Fermentas). For RT-PCR, 2 µl of cDNA was used per reaction. For quantitative real time PCR 1 µl of cDNA was used per reaction with 2x SYBR-Green Master Mix (Peqlab) on a Mastercycler ep realplex S (Eppendorf). *UBC* (AT5G25760) and *eIF4G* (AT3G60240) were used as reference genes. Primers are listed in Table S1. Triplicates of each sample were analyzed. The change of expression between two samples was determined by the 2-ΔΔCT Method according to (Livak & Schmittgen, 2001).

### Whole-mount in situ hybridization (WISH)

WISH on unfertilized pistils was carried out according to Hejatko et al. (2006). PCR fragments covering 313 bp of the 3’ region of *ENDOL7* (At1g79800) and 303 bp of the 3’ region of *AGO1* (At1g48410) were amplified using primer pairs ENODL7_F/ENODL7_R and AGO1_F/ AGO1_R, respectively, and cloned into pCR™II-TOPO (ThermoFisher). The coding sequences of *ASP* (At1g31450, 1338bp) and *SBT4.13* (At5g59120, 2196bp) were amplified from cDNA using primer pairs ASP_F/ASP_R and SBT4_F/SBT4_R, respectively, and cloned into pENTR/D-TOPO (Thermo Fisher Scientific). Primers are listed in Table S1. Probes were synthesized using the DIG RNA Labeling Mix kit (Roche, Product No. 11175025910) with linearized plasmid as a template. Probes for *SBT4.13* and *ASP* were fragmented to 300 bp. For small RNA detection by WISH, miRCURY LNA™ enhanced probes were ordered from EXIQON and 5’ DIG-labeled miRCURY LNA™ scramble-miR was used as negative control (Table S1). To prevent wash out of small RNAs from the sample an additional EDC-fixation step was carried out following company instructions (http://www.exiqon.com/ls/documents/scientific/edc-based-ish-protocol.pdf) and as described previously (Gosh Dastidar et al., 2016). For miRCURY LNA™ enhanced probes the hybridization and washing steps were carried out at 48°C with a probe concentration of 10-20 nM. Gene-specific probes were hybridized and washed at 55°C.

### Northern blots

Small RNAs were separated on 12% Polyacrylamid gels containing 7.5M Urea as previously described (Meister et al., 2004). After transfer to Hybond-N (Amersham Biosciences) and crosslinking with EDC (1-Ethyl-3-(3-dimethylaminopropyl) carbodiimide), membranes were probed with 5’ (γ-^32^P)ATP end-labeled antisense oligonucleotides directed against the miRNA of interest. Northern blots were stripped and re-probed for U6 snRNA as loading control. Oligonucleotide sequences are listed in Table S1.

### Construction of Reporter Transgenes

For promoter-reporter constructs, the respective promoter regions were amplified from genomic DNA using Phusion DNA Polymerase (New England Biolabs) and promoter-specific primers (Table S1) and cloned into pENTR/D-TOPO (Thermo Fisher Scientific) to generate entry clones. The promoters were sequenced and subsequently recombined into a pGREEN II-based vector containing a Gateway^®^ cassette in front of a nuclear localized 3xGFP sequence and a *NOS* transcriptional terminator (Takeda & Jürgens, 2007) using LR Clonase II (Thermo Fisher Scientific). Transgenic plants were generated by *Agrobacterium*-mediated transformation of inflorescences, after transferring the expression vectors into *Agrobacterium* GV3101 pMP90 pSOUP. Transgenic seedlings were selected with 200 mg/l glufosinate ammonium (BASTA^®^; Bayer Crop Science) supplemented with 0.1% Tween-3 to 5 days after germination. Pistils of transgenic plants were dissected as described (Sprunck et al., 2012) and analyzed for fluorescent reporter activity by confocal laser scanning microscopy (CSLM).

### Microscopy

Confocal laser scanning microscopy was performed on a Zeiss Axiovert 200M inverted microscope equipped with a confocal laser scanning module (LSM 510 META) and on an inverted Leica SP8 CLSM with HC PL APO CS2 2 40x/1.30 Oil and HC PL APO CS2 2 63x/1.30 glycerol objectives, respectively. The 488 nm-argon laser line was used for excitation of GFP and emission was detected with a BP 505-550 filter (LSM550 META) and from 500 to 535 nm by a hybrid detector (SP8), respectively. Cleared ovules were imaged with the Zeiss Axio Imager.M2 microscope with ApoTome.2 using a Plan-Apochromat 40x/1.4 NA oil DIC objective and differential interference contrast.

### Data accessibility

RNA-seq data have been deposited in NCBI GEO (https://www.ncbi.nlm.nih.gov/geo), accession number XXXXX. Arabidopsis egg cell RNA-seq data are also integrated in the open source web server CoNekT (Proost & Mutwil, 2018).

## Results

### Generation and characterization of RKD2-induced callus-like tissue

Köszegi et al. (2011) showed that ectopic expression of the egg cell-specific transcription factor RKD2 in Arabidopsis induced the formation of proliferating tissue that adopted transcriptome features of the egg cell. We aimed to utilize a similar RKD2-induced callus as a tool to elucidate small RNA pathway components in the Arabidopsis egg cell. To ectopically express an RKD2-GFP fusion protein in transgenic seedlings, we cloned *RKD2-GFP* under control of the *CaMV 35S* promoter (*35Sp:RKD2-GFP*). Proliferating cell masses occasionally appeared on the primary roots of transgenic seedlings around three weeks after germination on selection medium (Fig. 1A). Two independent RKD2-induced cell lines were propagated as calli on hormone-free selection medium (Fig. 1B). As a control tissue, we produced calli by cultivating root explants of Arabidopsis seedlings on callus-inducing medium (CIM) supplemented with auxin and cytokinin (Fig. 1C). Fluorescence of RKD2-GFP was regularly assessed in the RKD2-induced calli (Fig. 1D) and confocal laser scanning microscopy revealed that RKD2-GFP locates, like expected, to the nucleus (Fig. 1E). RT-PCRs were performed to detect transcripts of genes with known expression in the Arabidopsis egg cell, including *WOX2* (*WUSCHEL-related homeobox*) (Haecker et al., 2004), *ABI4* (*ABA INSENSITIVE 4*) (Wang et al., 2010), *SIAH* (*SEVEN IN ABSENTIA Homolog*) and *DRP4A* (*DYNAMIN RELATED PROTEIN 4A*) (Köszegi et al., 2011). Expression of *At5g01150* was tested as this gene shares sequence similarity to the egg cell- and proembryo-specific cDNA clone EC1-4 from wheat (Sprunck et al., 2005). The presence of known egg cell transcripts in the RKD2-induced callus suggested egg cell-like features, whereas these transcripts were not detectable in the auxin- and cytokinin-induced control callus (CIM) (Fig. 1F). Notably, the two RKD2-induced cell lines exhibited stable growth and RKD2-GFP expression throughout years of culture on hormone-free selection medium.

**Fig. 1.**
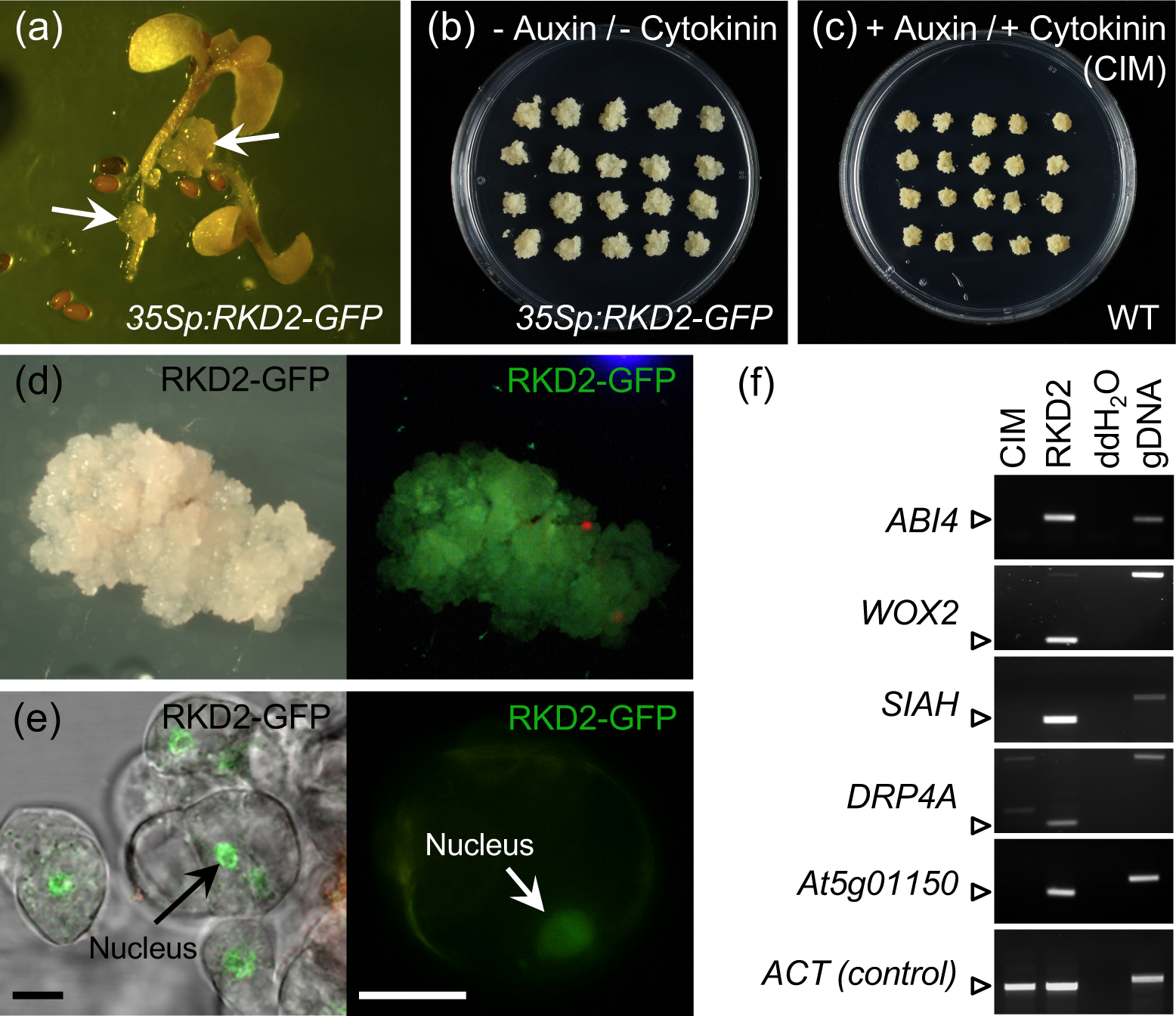
Generation of RKD2-induced egg cell-like callus. (a) Germinated Arabidopsis seedlings, transgenic for *35Sp:RKD2-GFP*. Callus-like cell masses (arrows) are visible at the primary roots of transgenic seedlings. (b) RKD2-induced cell line, propagated as calli on hormone-free medium. (c) Control calli, derived from seedling root segments cultured on callus induction medium (CIM) supplemented with auxin and cytokinin. (d) Bright field and fluorescence image of an egg cell-like callus expressing RKD2-GFP. (e) Cell cluster and single cell from RKD2-induced callus, with green fluorescence of RKD2-GFP in the nucleus. (f) Validation of egg cell-like cell fate by RT-PCR, using primer pairs specific for known egg cell-expressed genes. Abbreviations: *ABI4, ABA INSENSITIVE 4*; *WOX2, WUSCHEL-related homeobox 2; SIAH, SEVEN IN ABSENTIA Homolog; DRP4A, DYNAMIN RELATED PROTEIN 4A.* Size bars, 10 µm.

### RNA-Seq analysis of differentially expressed genes

To identify differentially expressed genes between the RKD2-induced callus and the CIM callus, three biological replicates for each callus type were used for RNA extraction, library preparation and Illumina Next-Generation Sequencing. After trimming, 52 to 63 million reads were retained per sample, of which between 93% and 96% were mapped by TopHat2 (Table S2). The total number of annotated genes in TAIR10.22 was 33,602, of which 24,928 remained after automatic independent filtering of low count genes by DESeq2. Pearson’s correlation coefficients indicated that the differences between the two callus types were considerably larger than between the three biological replicates (Fig. S1). 6,137 genes with a log2-fold threshold of 2 and a false-discovery rate (multiple testing adjusted p-value) smaller than 0.00001 were considered differentially expressed in the RKD2-induced callus when compared with the CIM callus (Table S3). Among those, 2,895 genes were induced (log2 FC ≥ 2) and 3,242 genes were repressed (log2 FC ≤ -2) in the RKD2 callus. Fig. 2a shows a Mean-versus-Average (MA) plot, providing an overview of gene expression differences depending on mean gene expression levels in the RKD2 and CIM calli. Red dots indicate significant differentially expressed genes (DEGs) and individually labeled genes were used for subsequent data validation.

**Fig. 2.**
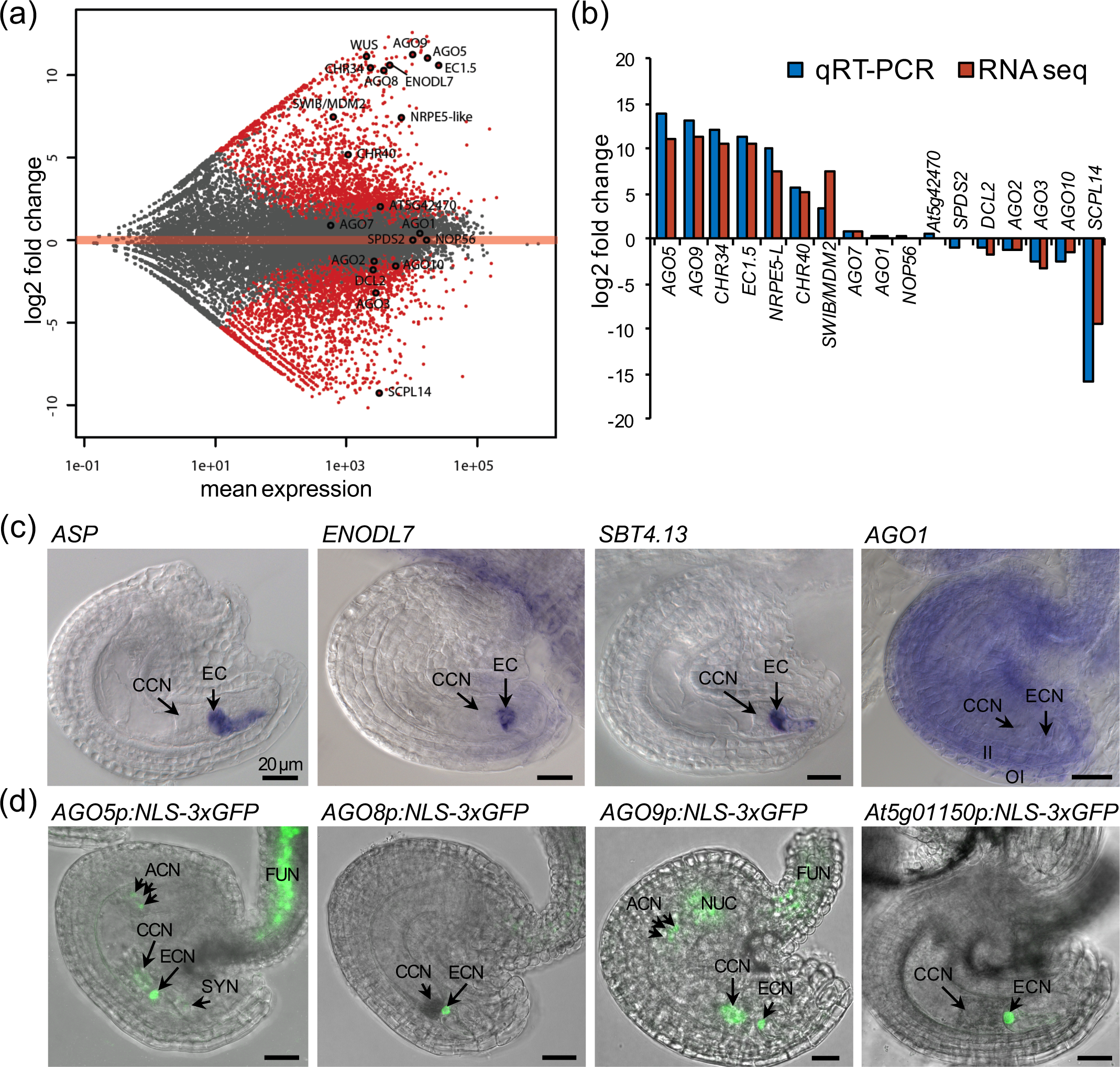
Analysis and validation of RNA-Seq data. (a) Mean-versus-Averages (MA) plot for differential expression analysis. Three RNA-Seq data sets for the RKD2-induced callus and the auxin-induced control callus (CIM), respectively, were analyzed. Each gene is represented with a dot. Genes with a False Discovery Rate (FDR) below 1*10-5 are shown in red. The horizontal axis shows the average expression over all samples; the Y axis indicates log2 fold changes between the RKD2-induced and the control callus. (b) Expression studies using real-time PCR, compared with RNA-Seq-based log2 fold changes. (c) Whole mount *in situ* hybridization using mature *Arabidopsis* ovules. Antisense probes are directed against mRNAs from RKD2-induced genes (*ASP, ENODL7, SBT4.13*) and *AGO1*, which is not differentially expressed. (d) Promoter-reporter studies. The nuclear localized GFP reporter (NLS-3xGFP) is expressed under control of promoters from genes with strong induction in the RKD2-induced callus. Merged fluorescence and brightfield CLSM images of mature ovules, prepared from transgenic plants, are shown. Abbreviations: ACN, antipodal cell nuclei; CCN, central cell nucleus; Chz, chalazal nucellus; EC, egg cell; ECN, egg cell nucleus; Fun, funiculus, II, inner integument; OI, outer integument. Size bars, 20 µm.

### Confirmation of differential expression between RKD2 and CIM calli by quantitative real-time RT-PCR analysis

Log2 fold changes from RNA-Seq data were validated by real-time RT-PCR using new cDNA batches from the two callus types, with *UBC21* (*UBIQUITIN-CONJUGATING ENZYME 21*) and *eIF4G* (*EUKARYOTIC TRANSLATION INITIATION FACTOR 4G*) as reference genes for normalization. We selected 17 genes covering the entire spectrum of log2 fold-changes in the RNA-Seq data and including genes with known roles in small RNA pathways (e.g., *AGOs, DCL, CHR40*), or with validated expression in the Arabidopsis egg cell (*EC1.5*) (Fig. 2b). RKD2-induced expression was validated for *AGO5* (*ARGONAUTE 5*), *AGO9* (*ARGONAUTE 9*), *CHR34* (*CHROMATIN REMODELING 34*), *CHR40* (*CHROMATIN REMODELING 40*), *ENODL7* (*EARLY NODULIN-Like 7*), *EC1.5* (*EGG CELL 1.5*), *NRPE5-L* (*NULEAR RNA POLYMERASE V subunit 5-Like*) and for the *SWIB/MDM2* family member At5g23480. Notably, the two chromatin-remodeling protein encoding genes *CHR34* and *CHR40* (also termed *CLASSY4*) were the only two family members with strong induction in the RKD2-induced callus, whereas others such as *CLASSY1* (*CHR38*), *BRM*/*CHR2* (*BRAHMA*), *PKL*/*CHR6* (*PICKLE*) and *SYD*/*CHR3* (*SPLAYED*) were repressed (Fig. S2a). Among all the plant-specific RNA polymerase Pol IV and Pol V subunits, *NRPE5-L* showed strongest induction in the RKD2-induced callus (Fig. S2b).

Likewise, significantly repressed genes in the RKD2-induced callus revealed a similar pattern in real-time RT-PCR, including *DCL2* (*DICER-LIKE 2*), the *ARGONAUTEs AGO2, AGO3* and *AGO10*, and the strongly repressed *SCPL14* (*SERINE CARBOXYPEPTIDASE-Like 14*). *AGO7* and *AGO1, NOP56* (*NUCLEOLAR PROTEIN 56*), the BRCA1-A complex subunit BRE-like (At5g42470) and *SPDS2* (*SPERMINE SYNTHASE 2*) were neither in RNA-Seq data nor in real-time RT-PCR differentially expressed (Fig. 2c).

### Expression of RKD2-induced genes in Arabidopsis ovules

RNA-Seq data were further validated by performing whole mount in situ hybridization (WISH) and promoter-reporter studies (Fig. 2c,d). WISH with mature Arabidopsis ovules revealed egg cell-specific expression of RKD2-induced *ASP* (*ASPARTYL PROTEASE*), *ENODL7* (log2FC of 10.6) and *SBT4.13* (*SUBTILASE 4.13*; log2FC of 3.37), whereas *AGO1* (log2FC of 0.41) showed a rather ubiquitous expression in the ovule (Fig. 2c). For promoter-reporter studies, a nuclear-localized 3xGFP reporter (NLS3xGFP) was expressed under control of selected promoter sequences in transgenic plants. At least five independent transgenic lines were investigated for GFP fluorescence by confocal laser scanning microscopy (Fig. 2d). Egg cell-specific reporter activity was observed when NLS3xGFP was expressed under control of the *AGO8* promoter (log2FC of 10.28) and the promoter of *At5g01150* (log2FC of 11.37), which encodes a Domain of Unknown Function 674 (DUF674) protein with sequence similarity to a wheat egg cell-expressed cDNA (Sprunck et al., 2005). The promoters of *AGO5* and *AGO9* revealed strong activity in the egg cell but were also active in the central cell, the antipodal cells and in the funiculus of mature ovules. The *AGO9* promoter was furthermore active in the chalazal nucellus of the ovule (Fig. 2d).

### Differentially expressed genes in the RKD2-induced callus and in egg cells

To investigate the gene expression profile of Arabidopsis egg cells, we isolated living cells from the Arabidopsis female gametophyte by micromanipulation as previously described (Englhart et al., 2017). We used the egg cell marker line EC1.1p:NLS3xGFP to be able to identify egg cells after they had been released from ovules treated with cell wall-degrading enzymes (Fig. 3a). Three replicates of 25 to 30 pooled egg cells each were used for RNA extraction, followed by library preparation from poly(A)+ RNA and Illumina Next-Generation Sequencing. After trimming, 47 to 57 million reads were obtained per replicate of pooled egg cells (Table S2) and 20,148 genes had a mean TPM of at least 1 (Table S4).

**Fig. 3.**
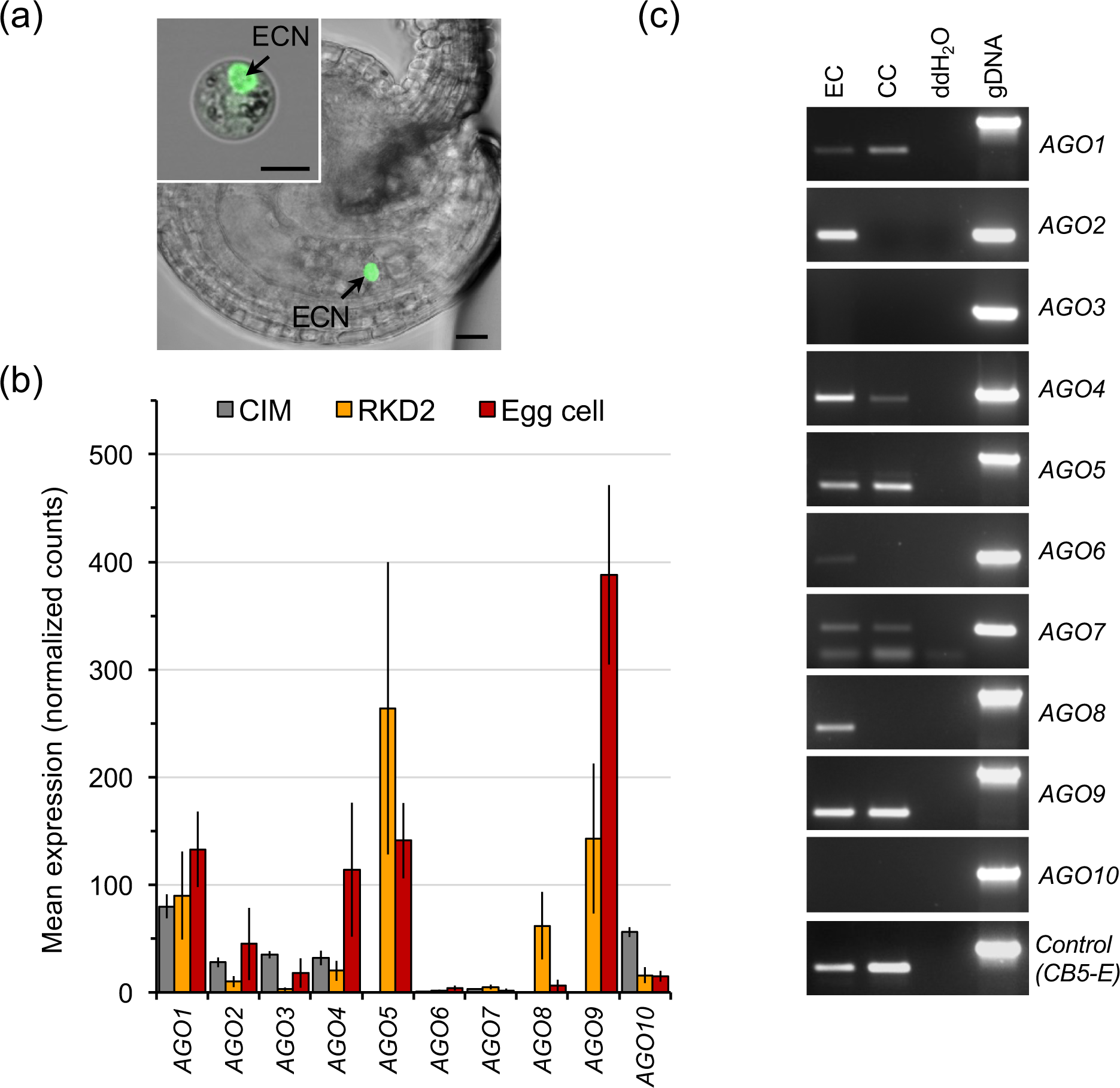
*ARGONAUTE* expression in egg cells. (a) Unfertilized ovule of Arabidopsis marker line EC1.1p:NLS3xGFP with green fluorescent egg cell nucleus (ECN). Inset in (a) shows an isolated egg cell of this marker line. Size bars, 10 µm. (b) Mean expression values for the ten Arabidopsis *AGOs* in the control callus (CIM), the RKD2-induced callus, and in egg cells. Expression levels are shown as average TPM (+/- SD). (c) RT-PCR with cDNA generated from each ten isolated Arabidopsis egg cells and central cells, respectively. Gene-specific *AGO* primers were used and *CB5-E* served as reference. Genomic DNA was amplified as positive control. Abbreviations: CC, central cell; EC, egg cell; ECN, egg cell nucleus; gDNA, genomic DNA.

Because of limited availability of egg cells, resulting in very low amounts of input RNA for sequencing, we had to use a different library preparation protocol than the one used for the CIM and RKD2-induced callus samples. To avoid artifacts when comparing sample groups, we performed differential expression analysis only for the comparison of RKD2-induced callus with CIM callus (Table S3). To compare the two callus types with the egg cell pools, we decided to not perform statistics because of the afore-mentioned issue, but only calculated transcripts per million (TPMs) to visualize differences in gene expression levels (Table S4).

In previous microarray-based expression studies, 99 genes were reported to be upregulated at least seven-fold in the RKD2-induced callus in comparison to seedlings and hormone-induced CIM callus, and were therefore considered as putative egg cell-specific genes (Kőszegi et al., 2011). We used these 99 genes to compare the three RNA-seq replicates of egg cells, RKD2-induced callus and hormone-induced CIM callus (Fig. S3). We furthermore added four *EC1* genes with known egg cell-specific expression to the analysis (Sprunck et al., 2012; Resentini et al., 2017), since only one *EC1* family member (*EC1.5*) was included. When we compared the gene expression profiles in the replicates generated from CIM and RKD2-induced calli, most TPMs were significantly higher in the RKD2-induced callus and therefore in line with the microarray-based expression data of Kőszegi et al. (2011). One exception was *CALS1* (*CALLOSE SYNTHASE 1*), which did not confirm the reported differential expression. However, the egg cell-specific genes *EC1.1, EC1.2, EC1.3* and *EC1.4* were not induced in the RKD2 callus. Furthermore, the embryo-expressed stem cell regulator *WUSCHEL* (*WUS*) and six other genes were RKD2-induced but not expressed in the egg cell RNA-seq libraries (Fig. S3). We therefore concluded that the RKD2-induced callus is not egg cell-like but rather adopts egg cell-related transcriptome features.

### Expression of small RNA pathway genes in the egg cell-related callus and in egg cells

ARGONAUTEs (AGOs) play a central role in RNA interference and microRNA pathways. When we compared mean expression values (average TPM, +/-SD) of the ten Arabidopsis *AGOs* we found *AGO5, AGO8* and *AGO9* to be specifically expressed both in egg cells and in egg cell-related RKD2 callus whereas they were absent in the CIM control callus (Fig. 3b), which is in line with our results obtained from real-time RT-PCR with the two callus tissues and promoter-reporter studies in Arabidopsis ovules (Fig. 2). Although the average TPM value for *AGO8* was very low, the presence of *AGO8* transcripts in egg cells was confirmed by RT-PCR (Fig. 3c). To investigate AGO expression in both female gametes we also included isolated central cells in our RT-PCR experiments. *AGO2, AGO6* and *AGO8* transcripts were only detected in egg cells, whereas amplification products for *AGO1, AGO4, AGO5, AGO7* and *AGO9* were detected in both female gametes (Fig. 3c). Inconclusive results were obtained for *AGO3* and *AGO10*, which were down-regulated in the egg cell-related callus and in egg cells (Fig. 2b; Fig. 4). Whereas no amplification products were obtained by RT-PCRs using isolated egg and central cells, egg cell RNA-seq data suggest presence of transcripts at least in the egg cell.

**Fig. 4.**
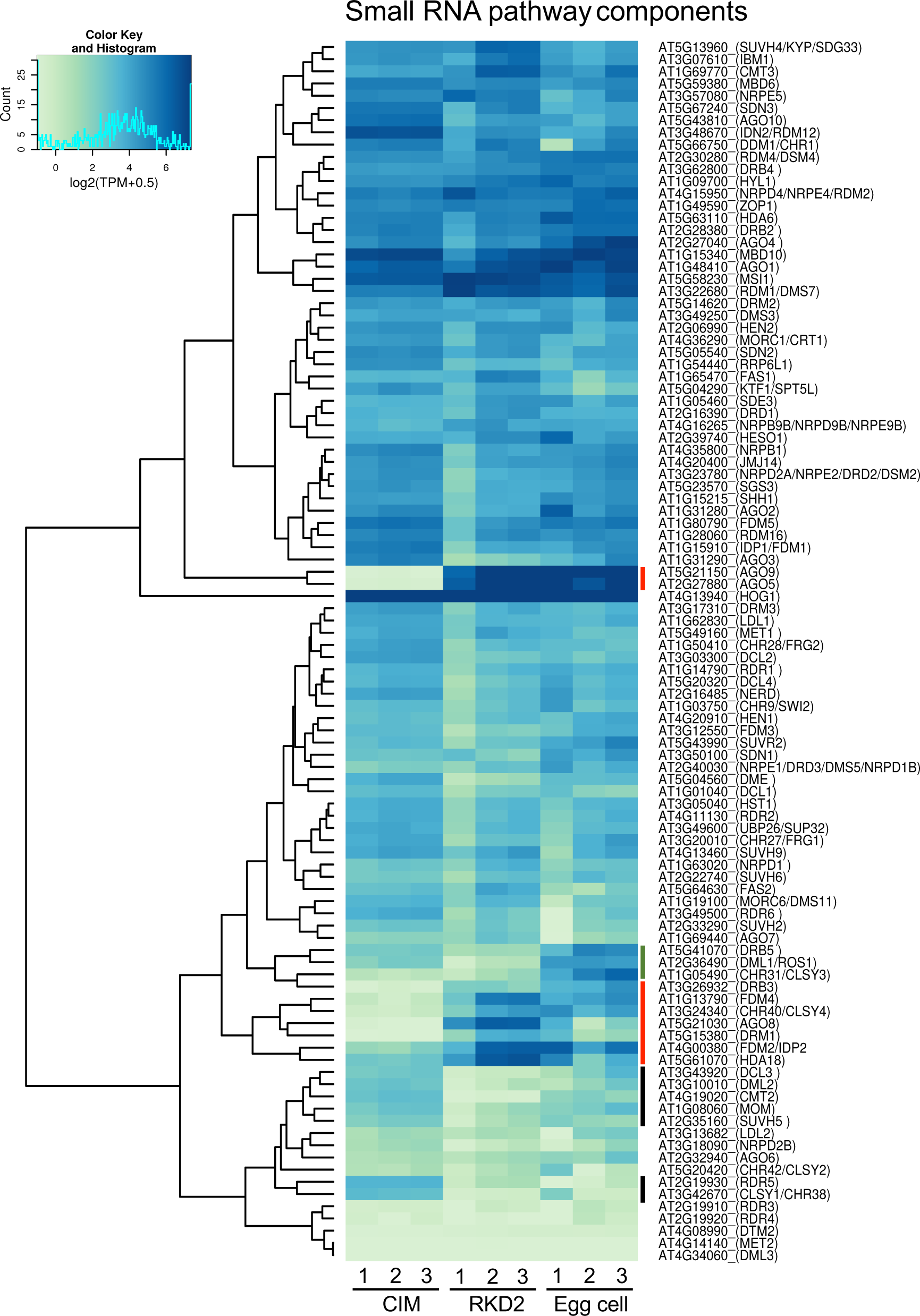
Expression pattern of genes known to be involved in small RNA pathways. The heatmap shows expression levels in the three biological replicates of hormone-induced control callus (CIM), RKD2-induced egg cell-like callus, and isolated egg cells. Color key indicates normalized and rlog transformed TPM values. Red bars label gene clusters with stronger expression levels both in egg cells and egg cell-like callus. Black bars label gene clusters with decreased expression in RKD2 callus and egg cells. Green bar labels genes with higher expression levels in egg cells but not RKD2 callus. Expression data of the genes shown in this ?gure are provided in Table S4.

A more global view on genes known to be involved in small RNA pathways and their differences in expression levels is shown in Fig. 4. Beside above mentioned *AGO5* and *AGO9*, the histone deacetylase *HDA18* and a cluster of genes comprising components of the RNA-directed DNA methylation (RdDM) pathway showed higher expression levels both in egg cells and the egg cell-related RKD2 callus, whereas expression levels were lower in the CIM callus. These are genes encoding DRB3 (DOUBLE-STRANDED RNA BINDING 3), the chromatin remodeling factor CHR40/CLASSY4, the SGS3-like dsRNA-binding proteins FDM2 and FDM4 (FACTOR OF DNA METHYLATION 2 and 4), and the *de novo* methyltransferase DRM1 (DOMAINS REARRANGED METHYLTRANSFERASE 1) controlling the maintenance of CHH methylation. By contrast, genes encoding the DNA glycosylase/lyase ROS1 (REPRESSOR OF SILENCING 1), DRB5 (DOUBLE-STRANDED RNA BINDING 5) and CHR31/ CLASSY3 showed stronger expression levels in egg cells but not in the RKD2-induced callus (Fig. 4).

### Expression of AGO proteins in the egg cell-related RKD2 callus

Stability and activity of small RNAs depend on AGO protein abundance. To test whether AGO expression in the egg cell-related callus is under translational control, we performed Western blot analyses with commercially available anti-AGO peptide antibodies and protein extracts from CIM callus, RKD2-induced callus and a set of sporophytic and reproductive tissues (leaf, root, flower and silique). AGO1 and AGO2 proteins were abundantly present in both callus types and detectable in all other tested tissues. AGO4 immunosignals were strongest in flowers and equally present in CIM and RKD2-induced callus. Much weaker signals were obtained for AGO6 in all tissues, including CIM and RKD2-induced callus. AGO5 and AGO9 proteins were neither detected in CIM callus nor in leaves and roots but abundantly present in flowers, siliques and in the RKD2-induced callus (Fig. 5), demonstrating that both transcripts are efficiently translated into proteins in the egg cell-related callus.

**Fig. 5.**
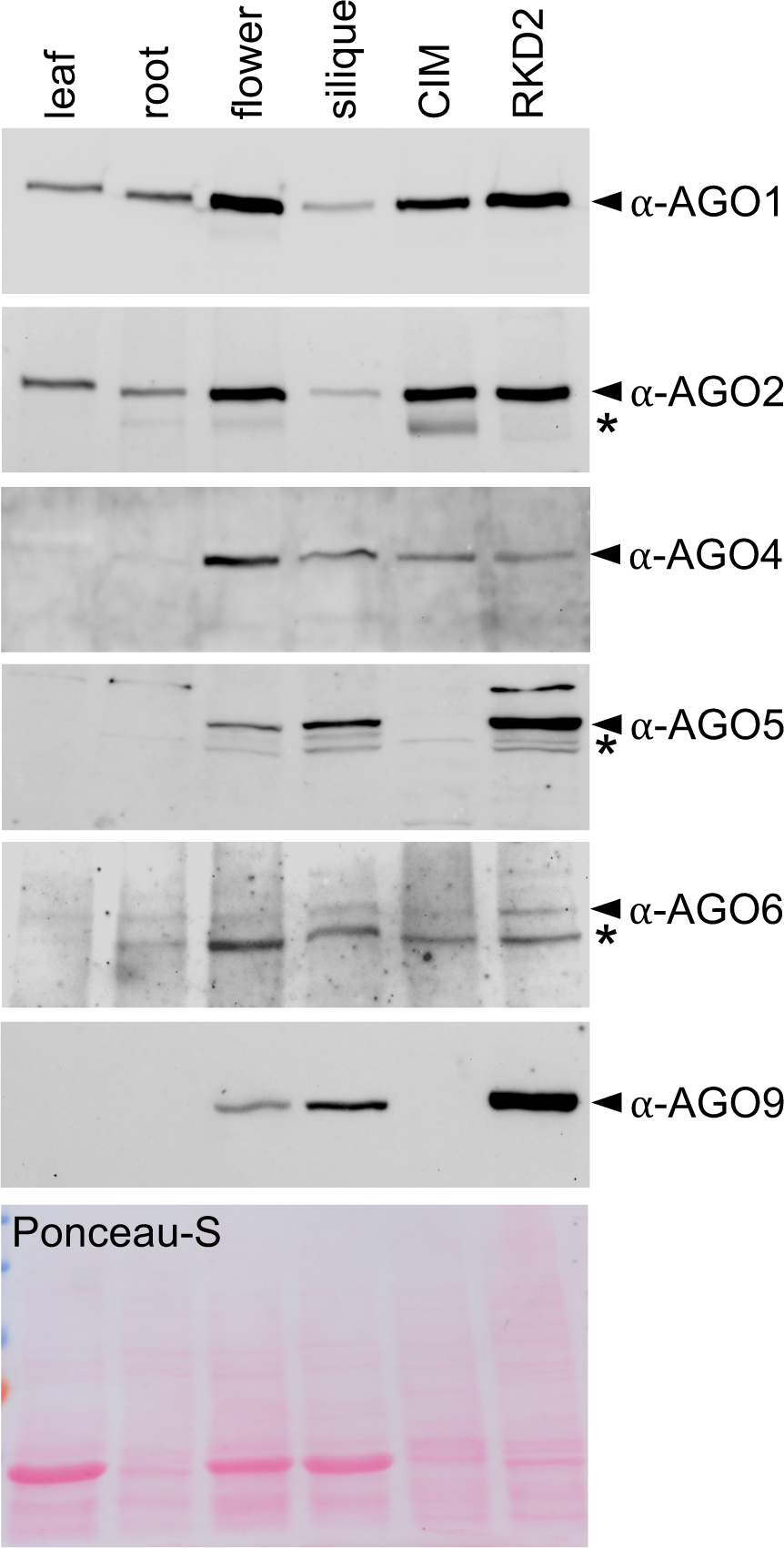
Western blot analyses showing ARGONAUTE (AGO) protein expression in the RKD2-induced egg cell-related callus, in the auxin-induced control callus (CIM), and in sporophytic and reproductive tissues of Arabidopsis. Six commercially available peptide antibodies were used, directed against AGO proteins as specified. Arrowheads label immunosignals matching with calculated and previously reported molecular weights. Asterisks mark putative degradation products. 20 μg total protein extract was loaded on each lane. Ponceau-S staining is shown as loading control.

### Small non-coding RNAs in the egg cell-related callus

Since we proved AGO protein abundance and egg cell-related transcriptome features, we considered the RKD2-induced callus suitable to investigate its small RNA (sRNA) profile in comparison to that of the hormone-induced CIM callus. We prepared six small RNA libraries (each three biological replicates) using the same batches of total RNA that had been used for mRNA-Seq for Illumina sequencing. After trimming, between 26 and 29 million small RNA (sRNA) reads were obtained for each sample of the hormone-treated CIM callus, whereas between 17 and 20 million sRNA reads were retrieved for each of the egg cell-related RKD2 callus samples (Table S2). For the CIM callus libraries, between 39 and 45% of the trimmed sRNA reads could be mapped uniquely to the Arabidopsis genome, and between 20 and 23% of the reads had multiple mapping locations. The libraries from the RKD2-induced callus contained between 8 and 12% reads which could be uniquely mapped, whereas between 30 and 38% of the reads had multiple mapping locations. The lower percentage of uniquely mapped reads in the sRNA libraries from RKD2 callus tissue was mainly caused by a significantly higher number of sRNAs derived from repetitive elements (e.g., transposable elements), which mapped to multiple genomic positions. Both uniquely and multi-mapping reads were considered for sRNA expression analyses.

We restricted further analyses to reads with length 18-24 nt, to exclude reads that did not derive from small RNAs but from degradation products of longer RNA specimen. This reduced the number of input reads to 10 to 14 million reads for the CIM callus samples and 4 to 5 million reads for the RKD2 callus samples. Mapping rates were higher for this subset, with 48-49% unique mappings for CIM callus samples and 17-21% for RKD2 callus samples, respectively, and 36-37% multi-mapping reads for CIM callus samples, and 70-75% multi-mapping reads for RKD2 callus samples (Table S2).

To compare the relative abundance of read length across sRNA feature types, we calculated relative read counts (percentages) for each sample, and summarized biological replicates by taking the median. The resulting profile of read length revealed that the majority of reads was either 21 nt or 24 nt long (Fig. 6a), which is in agreement with the miRNA and siRNA classes of small RNAs in Arabidopsis. However, 24-nt sRNAs were most abundant in the RKD2-induced callus, whereas the highest percentage of counts in the CIM callus comprised 21-nt sRNAs, likely reflecting stronger tendencies towards post-transcriptional silencing in the CIM callus and RNA-dependent DNA methylation in the egg cell-related callus, respectively. To functionally dissect the reads with specific lengths, we aligned them to the genomic features annotated in the TAIR database (arabidopsis.org). Notably, the egg cell-related callus produced a proportionately higher number of transposable element (TE) sRNA counts (47% of all counts, versus 20% in CIM callus). TE-derived siRNAs were prominent among the 24 nt reads but also among the 21-nt and 22-nt long reads, indicating the posttranscriptional degradation of transposable elements activated in the egg cell-related callus.

**Fig. 6.**
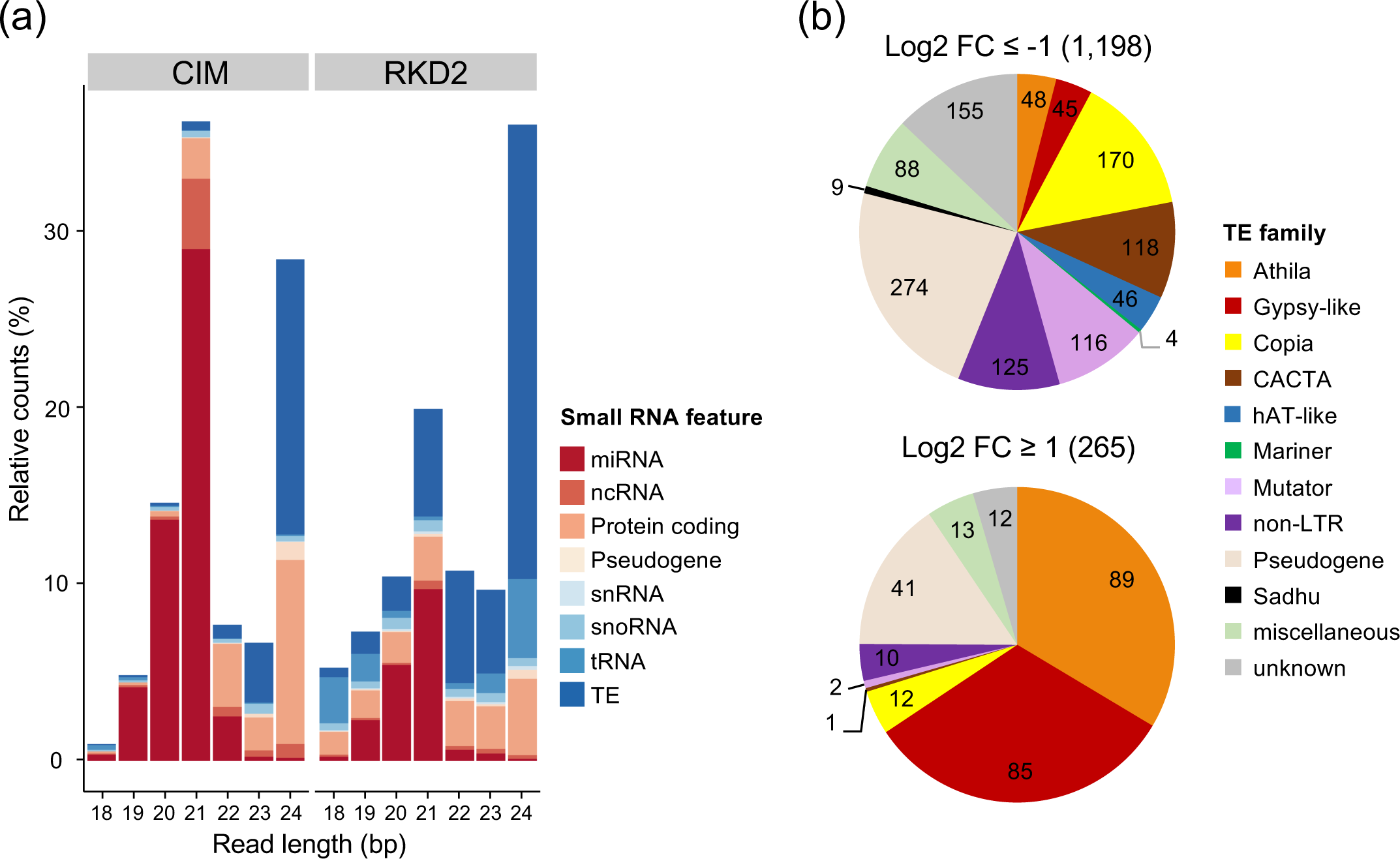
Differentially expressed small noncoding RNAs in the two distinct callus types, analyzed by small RNA sequencing. (a) Distribution of small RNA reads from RKD2 and CIM calli into different lengths and genomic features. Small RNA reads were classified by length and alignment position within genomic features as annotated in TAIR10. Genomic features comprise microRNA (miRNA), non-coding RNA (ncRNA), protein coding gene, pseudogene, small nuclear RNA (snRNA), small nucleolar RNA (snoRNA), transfer RNA (tRNAs) and transposable element (TE). For each read length, the read distribution into genomic features was calculated relative to all reads ranging from 18 to 24 nt and aligning to any of the features mentioned above. Note the decreased percentages of reads aligning to *miRNA* loci and the increased percentages of 20 to 24 bp reads aligning to *TE* loci in the RKD2-induced callus. (b) Differentially expressed *TE* genes with mapped small RNA reads in the RKD2 callus and their distribution across TE families. Numbers correspond to genes with mapped small RNA reads which were either induced (log2 FC ≥1; P < 0.0001) or repressed (log2 FC ≤-1; P < 0.0001) in the RKD2-induced callus in comparison with the CIM callus. Genes annotated as transposable_element_gene were grouped in miscellaneous, unknown, or pseudogene TEs according to additional informations. Numbers in brackets represent total number of TE gene loci.

**Fig. 7.**
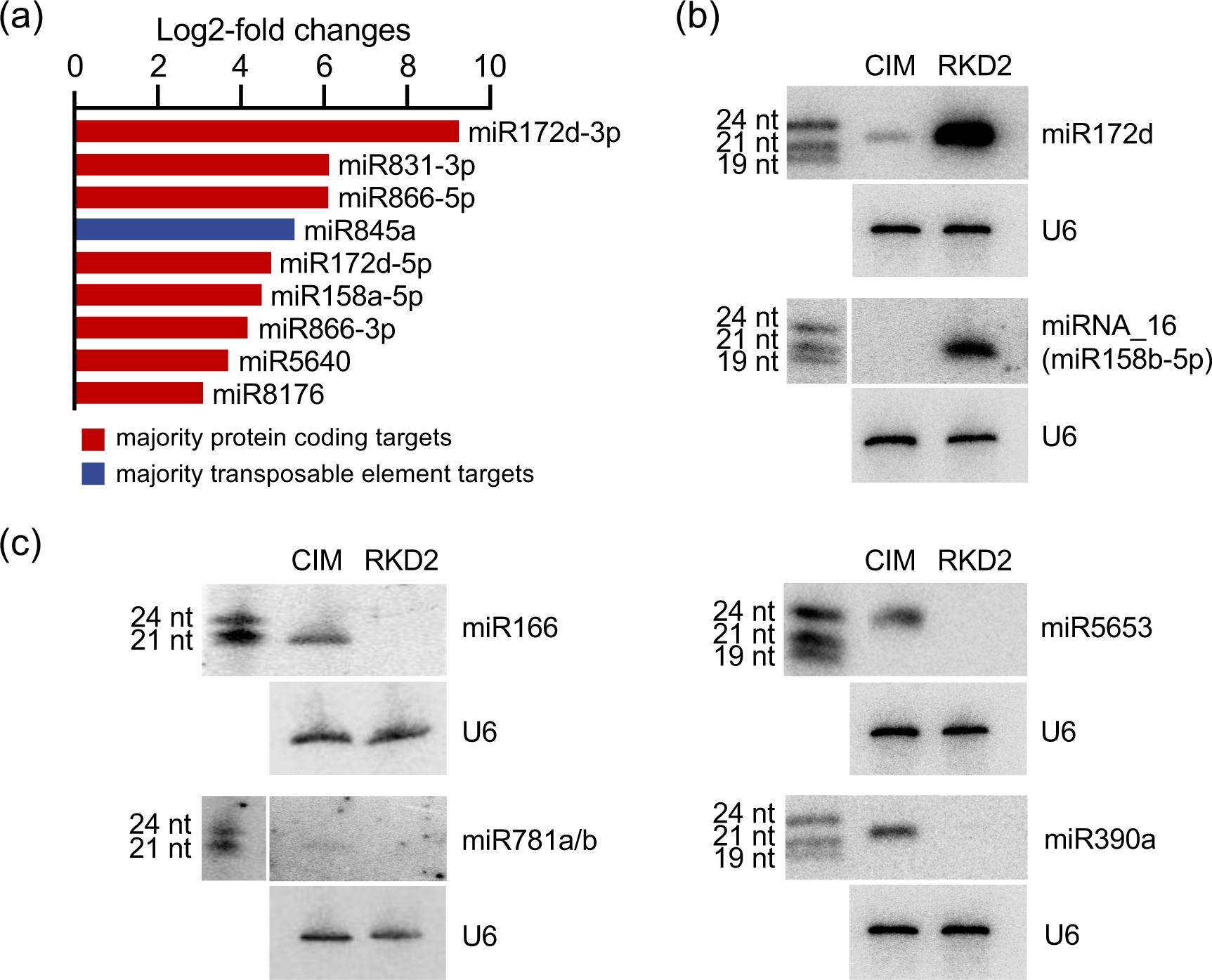
Differentially expressed miRNAs. (a) Log2-fold changes of induced miRNAs in the RKD2 egg cell-like callus. Bar colors are based on miRNA targets being mostly protein-coding genes (red) or transposable elements (blue) in plant small RNA target analysis (psRNATarget). Repressed miRNAs are shown in Fig. S4. (b), (c) Validation of differentially expressed miRNAs by Northern blot hybridization. Total RNA from RKD2-induced callus (RKD2) and hormone-induced control callus (CIM) was used for hybridization with probes directed against miRNAs with induced (b) or repressed (c) expression in the RKD2 callus. Northern blots were stripped and reprobed for U6 snRNA as loading control. The position of RNA size markers, electrophoresed on the same gel, is shown to the left of the blots. Log2-fold changes of repressed miRNAs in the RKD2 egg cell-like callus are shown in Fig. S4.

Increased levels were furthermore obtained for 18, 19, 23 and 24 nt long reads originating from *tRNA* loci (10.8% of all counts in RKD2 callus, versus 0.7% in CIM callus) (Fig. 6a). On the other hand, the egg cell-related callus revealed reduced levels of miRNAs (19% of total reads versus 51% in CIM callus). A considerable fraction of reads with length 21 nt and longer aligned to protein-coding genes in the CIM callus, whereas in the RKD2 callus reads mapping to protein-coding regions were represented across the complete range of read lengths analyzed. However, the total number of reads aligned to protein-coding regions did not differ much between the two types of calli (18% in CIM callus versus 16% in RKD2 callus). These reads can stem from siRNAs targeted against protein-coding genes and from degradation products of protein-coding genes.

### Expression changes of TE sRNAs and miRNAs in the egg cell-related RKD2 callus

For differential expression analysis of small RNAs with DESeq2, we considered small RNA-seq reads of lengths 18 to 24 nt mapping to a genomic feature annotated in TAIR10.22 for library size normalization and focused on miRNAs and transposable element (TE)-derived small RNAs.

#### Small RNAs silencing transposable elements

Small RNA read counts aligning to the antisense strand of TEs were investigated in more detail for their differential expression and distribution across TE families. Among the 1,463 *TE* loci with mapped sRNA reads, the majority (81.9%) was repressed in the RKD2 callus, suggesting that the corresponding siRNAs mediate transcriptional silencing to prevent deleterious reactivation of transposons (Fig.6b and Table S5). The largest groups among the repressed *TE* were formed by *copia-like* long terminal repeat (LTR) retrotransposons and non-LTR retrotransposons (LINEs), followed by class II DNA transposons from the *CACTA, hAT* (*hobo/Ac/Tam3*)-like and *Mutator* superfamilies. Nevertheless, we also detected 265 transcriptionally active *TE* loci with matching sRNA reads in the egg cell-related callus, matching with previous observations that reactivation of TEs occurs specifically during female and male gamete formation (Slotkin et al. 2009; Olmedo-Monfil et al., 2010). Notably, the majority (66%) of these induced *TE* loci belong to the *Gypsy-like* superfamily of long terminal repeat (LTR) retrotransposons and more than half of those were *Athila* LTRs (Fig.6b).

#### miRNAs

To quantify the differential expression of annotated and novel miRNAs, we substituted the miRNAs annotated in TAIR10.22 with Arabidopsis miRNAs from miRBase and predicted novel miRNAs using the miR-PREFeR prediction tool (Lei and Sun, 2014) before running DESeq2. Those miRNAs with annotated sequences in the miRBase sequence database (Kozomara and Griffiths-Jones, 2014) were referred to as “known”. Differential expression profiling of known miRNAs revealed clear differences between the egg cell-related RKD2 callus and the hormone-induced CIM callus. In total, 96 known miRNAs were differentially regulated with a log2 fold change of at least 2 (Table S6). Notably, the great majority of these miRNAs (88) showed reduced abundance in the RKD2 callus when compared with the control callus (Fig. S4), whereas only 9 miRNAs were induced. Considering -3p and -5p mature forms separately, the strongest induced miRNA in the RKD2 callus was miR172d-3p (log2FC of 9.2), followed by miR831-3p and miR866-5p (both log2FC of 6.1) (Fig.7a). Highest read counts in the RKD2 callus were derived for upregulated miR158a-5p (average counts 6,252; log2FC of 4.5), miR866 (average counts 1,549; log2FC of 6.1), and miR5640 (average counts 1,615; log2FC of 3.7) (Table S6).

The strongest repressed miRNA was miR165b (log2FC of -14.52) with an average of 7,098 counts in the CIM callus but no counts in the RKD2 callus. The most abundant miRNAs in the CIM callus were those derived from *MIR165* and *MIR166* loci (miR165a,b and 166a,c,f,g). Furthermore, miR157a,b,c (log2FC of -9.8 to -5.1), miR156a,c,d,e (log2FC of -9.4 to -3.7) and miR8167 (log2FC of -9.2 to -4.8) were among the strongest repressed miRNA in the CIM callus (Fig. S4 and Table S6).

Beside known miRNAs we identified 30 potential novel miRNAs which were differentially expressed. Comparisons of the genomic coordinates of these “novel” miRNAs with genomic coordinates of miRNAs in miRBase revealed that some of them correspond to stem-loop sequences of known MIR precursors but differ from known miRNAs derived from these stem loops. In total, 20 differentially expressed novel miRNAs were linked to known MIR precursors, whereas 10 differentially expressed novel miRNAs mapped to new gene loci (Table S7). The differential expression of six selected known and novel miRNAs was successfully validated by Northern blot hybridization (Fig.7b).

### Predicted and experimentally validated targets of differentially expressed miRNAs

The identification of target genes of the differentially expressed miRNAs was performed using the plant small RNA target analysis tool psRNAtarget. We then analyzed the differential expression of predicted targets in the RKD2-induced callus, particularly taking account of miRNA-targeted cleavage which should result in negatively correlated expression tendencies of the miRNA targets. Table 1 shows induced miRNAs in the egg cell-related callus, their predicted or experimentally validated targets and the changes in target gene expression in the RKD2-induced callus. Six *Arabidopsis* APETALA2-LIKE (AP2-like) transcription factors are known targets for miR172 (Aukerman and Sakai, 2003; Schmid et al., 2003; Mathieu et al, 2009). Two of them, SMZ (*SCHLAFMUTZE*) and *TOE3* (*TARGET OF EARLY ACTIVATION 3*), are validated targets of miR172d and their downregulation in the RKD2 callus suggests miR172d-induced cleavage of transcripts. Notably, we found miR845a to be upregulated in the RKD2 callus, although the miR845 family was reported to be preferentially expressed in mature *Arabidopsis* pollen (Creasey et al., 2014; Borges et al., 2018). The *Gypsy*-like retrotransposon *Athila* (AT1G43060) is a predicted target for miR845a-guided cleavage and its expression is downregulated in the RKD2 callus. Opposing expression tendencies were also obtained for targets predicted for the other seven RKD2-induced miRNAs (Table 1), and for targets of miRNAs which were strongly repressed in the RKD2 callus (Table S8).

**Table 1.** Target prediction for differentially upregulated miRNAs and target gene expression in the RKD2-induced callus.

| miRNA name | log2FC miRNA<sup>1</sup> | target accession | log2FC target | p/v<sup>2</sup> | Inhibition | target symbol | target name / target family |
| --- | --- | --- | --- | --- | --- | --- | --- |
| miR172d-3p | 9.2 | AT3G54990 | -4.2 | v | cleavage | SMZ | SCHLAFMUTZE/Apetala2-like transcription factor |
|  |  | AT5G67180 | -2.9 | v | cleavage | TOE3 | Target of early activation tagged (EAT) 3/Apetala2-like transcription factor |
|  |  | AT3G47360 | -3.4 | p | cleavage | HSD3 | Hydroxysteroid dehydrogenase 3 |
|  |  | AT5G12900 | -2.5 | p | cleavage |  | DNA double-strand break repair RAD50 ATPase |
| miR831-3p | 6.1 | AT1G69220 | -1.7 | p | cleavage | SIK1 | Serine/threonine kinase 1 (Hippo/STE20 homolog) |
|  |  | AT5G10320 | -3.4 | p | cleavage |  | ATP synthase subunit B |
|  |  | AT1G63100 | -3.4 | p | cleavage |  | GRAS family transcription factor |
|  |  | AT2G43445 | -4.2 | p | cleavage |  | F-box and associated interaction domains-containing protein |
| miR866-5p | 6.1 | AT5G06510 | -3.0 | p | cleavage | NF-YA10 | Nuclear Factor Y, subunit A10 |
| miR845a | 5.3 | AT1G43060 | -2.7 | p | cleavage |  | gypsy-like retrotransposon family (Athila) |
| miR172d-5p | 4.7 | AT1G70782.1 | -2.1 | p | cleavage | CPuORF28 | Conserved Peptide upstream Open Reading Frame 28 |
|  |  | AT3G23810 | -3.0 | p | cleavage | SAHH2 | S-adenosyl-L-homocysteine (SAH) hydrolase 2 |
| miR158a-5p | 4.5 | AT1G79680 | -3.9 | p | translation | WAKL10 | WALL ASSOCIATED KINASE-LIKE 10 |
|  |  | AT1G55970 | -1.7 | p | cleavage | HAC4 | Histone acetyltransferase of the CBP family 4 |
| miR866-3p | 4.2 | AT3G05380 | -1.9 | p | cleavage | ALY2 | ALWAYS EARLY 2 Myb-DNA binding factor |
|  |  | AT4G20110 | -2.4 | p | cleavage | VSR7 | VACUOLAR SORTING RECEPTOR 7 |
|  |  | AT5G18370 | -1.7 | p | cleavage | DSC2 | DOMINANT SUPPRESSOR OF CAMTA3 NUMBER 2 |
|  |  | AT1G43886 | -2.5 | p | cleavage |  | copia-like retrotransposon family |
| miR5640 | 3.7 | AT1G61850 | -1.1 | p | cleavage |  | Non-specific lipase |
| miR8176 | 3.1 | AT3G20935 | -5.9 | p | cleavage | CYP705A28 | Cytochrome P450, family 705, subfamily A, polypeptide 28 |
|  |  | AT4G37400 | -6.9 | p | cleavage | CYP81F3 | Cytochrome P450, family 81, subfamily F, polypeptide 3 |
|  |  | AT1G15160 | -2.3 | p | cleavage |  | MATE efflux family protein |

Only miRNAs and targets with log2-fold changes of $\geq 2$ are shown
<sup>1</sup> log2 fold change of miRNA expression (RKD2-induced vs. CIM callus)
<sup>2</sup> p, predicted target; v, validated target. Target predictions are based on psRNAtarget (Dai and Zhao, 2011). Validated targets according to miRTarBase 7.0 (http://mirtarbase.mbc.nctu.edu.tw/php/download.php) and Dai et al. (2018). The complete list of differentially-expressed miRNAs and their predicted/validated targets is given in Table S8.

Examples are targets for miR5653 (log2FC of -9.6), miR319b (logFC of -8.4), miR159b (logFC of -7.0), and miR781a,b (log2FC -2.4). Remarkably, miR5653 is predicted to target *ABI4* (*ABA INSENSITIVE 4*), which is strongly induced in the RKD2 callus (log2FC of 10.5; Fig. 2b) and shows egg cell-specific expression in mature ovules (Wang et al., 2010). The miR319 family is known to target *TCP* transcription factors and one validated target for miR319b is *TCP4*, exhibiting elevated expression in the RKD2 callus (log2FC of 4.4). Likewise, a known target for miR159b (*MYB101*) shows opposing expression in the RKD2 callus (log2FC of 4.1). One predicted target of miR781a,b encodes a SWIB/MDM2, Plus-3 and GYF domain-containing protein (At5g23480). Notably, At5g23480 is egg cell-expressed and strongly induced in the RKD2 callus (log2FC of 9.74), which was verified by real time RT-PCR (Fig. 2b). A comprehensive overview on differentially expressed known and novel miRNAs, together with target predictions by psRNAtarget is given in Tables S8 to S10.

### Differentially expressed small non-coding RNAs in the female gametophyte

We performed small RNA whole mount *in situ* hybridization (WISH) with DIG-labeled locked nucleic acid (LNA) antisense probes to examine the spatial distribution of differentially expressed miRNAs in Arabidopsis Col-0 ovules (Fig. 8). In the sporophytic tissues of the ovule, RKD2-induced miR172c,d was detected in the inner integuments and the chalazal nucellus but not in the outer integuments. Within the female gametophyte, signals for miR172c,d were detectable in the nucleus and cytoplasm of egg cells, central cells, and synergid cells. A different pattern of expression was obtained for the other miRNAs. Whereas signals for RKD2-induced miR8176 especially accumulated in the nucleoli of egg cells, synergid cells and central cells, miR5640 was detected in the nucleus and cytoplasm of the egg cell and the two synergid cells. Faint miR5640 signals were also detectable in the nucleus of the central cell. Within the female gametophyte, novel miR_95 (log2FC of 1.6 in the RKD2-induced callus) was detected in the egg cell, the two synergid cells and the central cells. Furthermore, novel miR_95 was detectable in sporophytic cells, specifically in inner and outer integument cells forming the micropyle of the ovule.

**Fig. 8.**
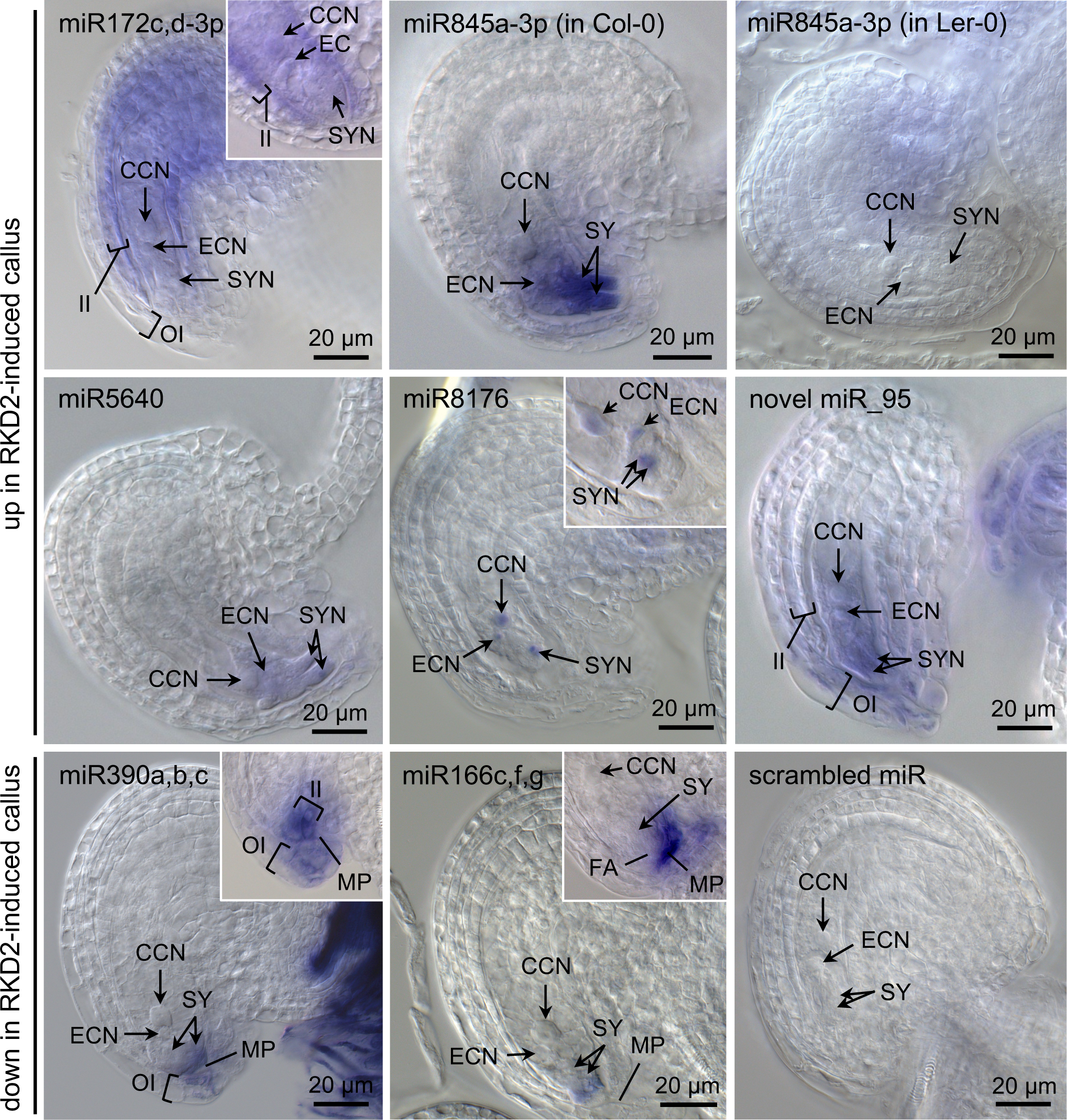
Localization of differentially expressed miRNAs in Arabidopsis ovules by whole mount in situ hybridization (WISH). MirCURY LNA™ detection probes were used for hybridization. When a probe can detect more than one miRNA isoform, this is indicated by the name (e.g., miR172c,d-3p). Insets show the micropylar region of another representative ovule hybridized with the same probe. WISH with scrambled miR, performed in parallel, served as control. Note that miR845a-3p is not detected in the female gametophyte of ecotype Landsberg erecta (Ler-0), which has a 1kb deletion at the *MIR845a* locus. All other ovules are of ecotype Columbia (Col-0). Abbreviations: CCN, central cell nucleus; EC, egg cell; ECN, egg cell nucleus; FA, filiform apparatus; II, inner integument; MP, micropyle; OI, outer integument; SY, synergid cell; SYN, synergid cell nucleus. Size bars, 20 μm.

Surprisingly, the RKD2-induced miR845a-3p, previously reported to be preferentially expressed in pollen (Borges et al., 2018), showed very strong signals in the two synergid cells, weaker signals in the egg cell and either no, or very faint signals in the central cell. Since the *Arabidopsis* ecotype Landsberg erecta (Ler-0) has a 1kb deletion at the *MIR845a* locus (Borges et al., 2018), we concomitantly performed WISH with ovules from this ecotype. We did not detect any signals in the cells of the Ler-0 female gametophyte, arguing for the specificity of the probe directed against miR845a-3p (Fig. 8).

We also included WISH experiments with miRNAs which were repressed in the RKD2 callus (Fig. 8; up in CIM callus). Indeed, the isoforms of miR390 and miR166, both strongly repressed in the RKD2-induced callus (log2FC of −8.9 to −6.0 for miR390b,c and logFC of −9.7 to −9.1 for miR166c,f,g, respectively), have not been detected in egg cells. However, these miRNAs showed a synergid-specific expression within the female gametophyte and exhibited a polar distribution towards their micropylar pole. Notably, we also detected signals of miR390b,c and miR166c,f,g in the micropyle of the ovule, suggesting them to be secreted by the synergid cells.

## Discussion

Members of the plant-specific RWP-RK DOMAIN CONTAINING (RKD) family of transcription factors have a conserved role as egg cell determinants in Arabidopsis and *Marchantia polymorpha* (Köszegi et al., 2011; Koi et al., 2016; Rövekamp et al., 2016). In Arabidopsis, five *RKD* genes are expressed during distinct stages of embryo sac development, with *RKD1* and *RKD2* being highly expressed in egg cells (Tedeschi et al., 2017). RKD2 was reported to induce cell proliferation and the activation of egg-like transcriptomic features when ectopically expressed in seedlings, protoplasts, or in sporophytic cells of the ovule (Köszegi et al., 2011; Lawit et al., 2013). We analyzed the transcriptome of own RKD2-induced cell lines in comparison to that of a hormone-induced CIM callus and found, with one exception, all putative egg cell-specific genes that have been reported by Kőszegi et al. (2011) to be differentially upregulated in the RKD2-induced callus.

We complemented the transcriptome data with RNA-Seq data generated from isolated Arabidopsis egg cells and found a largely overlapping expression pattern of genes which are induced in both the egg cell and the RDK2 callus, including *AGOs* and known or, so far, unexplored chromatin regulators. These include, for example, histone deacetylase HDA18 which might act, like HDA6, in RNA directed DNA methylation, the chromatin remodeling factors CHR34 and CHR40/CLASSY4, a *SWIB/MDM2* family member encoded by At5g23480 and the *POL V subunit 5-like* protein which might all fulfill specific functions in the egg cell. Nevertheless, we also found differences between the RKD2-induced callus and the egg cell transcriptome since, for example the embryo-expressed stem cell regulator *WUSCHEL* (*WUS*) is induced in the RKD2 callus although not expressed in the egg cell, whereas four of the five egg cell-specific *EC1* genes were not RKD2-induced. These variations in expression suggest that certain aspects of the egg cell transcriptome are activated in the RKD2-induced callus rather than all transcriptome features of the egg cell.

Gamete development is controlled by unique gene expression programs and involves epigenetic reprogramming of histone modifications and DNA methylation (Jullien & Berger, 2010; Baroux et al., 2011). However, demethylation and removal of repressive histone marks render the genome particularly susceptible to increased TE mobilization. In animals, the PIWI/piRNA pathway targets reactivated TEs post-transcriptionally and through the induction of epigenetic changes at the loci from which they are expressed (Russell & LaMarre, 2018). Although flowering plant genomes lack the germline-expressed Piwi subfamily of Argonautes and piRNAs, certain plant AGOs are preferentially expressed in their reproductive cell lineages (Grimson et al., 2008; Borges and Martienssen, 2015). In rice, mutations in the AGO5 homolog *MEIOSIS ARRESTED AT LEPTOTENE 1* (*MEL1*) induce precocious meiotic arrest and male sterility, due to defects in large-scale meiotic chromosome reprogramming and histone hypermethylation (Nonomura et al., 2007; Liu et al., 2016). Maize AGO9 is expressed in ovule somatic cells surrounding the megaspore mother cell (female meiocyte) and contributes to non-CG DNA methylation in heterochromatin. Furthermore, chromosome segregation is arrested during meiosis in maize *ago9* mutants (Singh et al., 2011). In Arabidopsis, at least two small RNA-mediated silencing pathways appear to act in somatic cells flanking the megaspore mother cell and the functional megaspore (haploid product of meiosis) to regulate female reproductive development. One pathway depends on AGO9 and 24 nt siRNAs to prevent sub-epidermal somatic cells from adopting megaspore-like cell identity (Olmedo-Monfil et al., 2010), whereas a second, independent small RNA-mediated pathway acts in the functional megaspore to promote the transition to megagametogenesis (Tucker et al., 2012).

However, *ago9* mutants of maize and Arabidopsis imply divergent mechanisms: Arabidopsis AGO9 represses meiocyte cell fate in somatic cells (Olmedo-Monfil et al., 2010), while maize AGO9 inhibits somatic cell fate in meiocytes (Singh et al., 2011). Moreover, the impact of small noncoding RNAs on chromatin changes, the transcriptional landscape and cell identity of egg cells is not yet explored. Our small RNA-seq data revealed that the egg cell-related callus is highly enriched in TE-derived small RNAs, whereas the ratio of miRNAs is decreased when compared with the hormone-induced CIM callus. Most of the TE loci aligning with the small RNA reads are downregulated. However, 265 TEs with mapped reads were transcriptionally active and they mainly belong to the *Gypsy-like/Athila* superfamily of LTRs. It will remain an interesting task for the future to identify genic mRNAs targeted by the TE-derived siRNAs, as they might be involved in regulating egg cell fate. Furthermore, AGO5 and AGO9 are translated into proteins in the RKD2-induced callus. The egg cell-related callus therefore represents a unique tool to overcome cell material limitations for protein biochemistry, which will allow the purification and analysis of bound small RNAs in an egg cell-related background.

During Arabidopsis male gametophyte development, several TEs were reported to be less methylated in the pollen vegetative cell due to reduced expression of DDM1 (DECREASE IN DNA METHYLATION1), and, thus, transiently reactivated. As consequence, miRNA-mediated post-transcriptional gene silencing of reactivated TEs generates 21-nt epigenetically expressed siRNAs (easiRNAs), which are transferred from the vegetative cell nucleus to the sperm cells to reinforce silencing of complementary TEs and to protect the male gametes (Slotkin et al., 2009; Creasey et al. 2014). In the female gametophyte, a similar scenario was proposed based on the observation that the central cell undergoes active DNA demethylation before fertilization, resulting in TE reactivation. Small RNA movement is also assumed between the central and the egg cell, based on the observation that an artificial microRNA, expressed in the central cell, targets cleavage of green fluorescent protein (GFP) RNA expressed in the egg cell (Ibarra et al., 2012).

Notably, our results from small RNA WISH with Arabidopsis ovules showed that most of the RKD2-induced miRNAs were also detectable in egg cell. One exception was miR158a-5p, which could not be detected in the ovule whereas signals were obtained in cells of the RKD2 callus (not shown). For the other miRNAs, signals were either detectable in egg cells, synergids and central cells, or in egg cells and synergids. Of special interest was miR845a which was reported to be preferentially pollen expressed and to target transposable elements (Borges et al., 2018). WISH signals for miR845a were much stronger in the synergid cells than in egg cells and could indicate that *MIR845a* is strongly expressed in the accessory synergid cells and moved to the egg cell to silence reactivated transposable elements in the female gamete. Intercellular communication of the synergid cells with the egg cell via small RNAs has not yet been reported and will be interesting to study in more detail in the future.

On the other hand, two CIM-induced miRNAs (miR166 and miR390) were detectable in synergid cells but not in egg cells. Interestingly, these miRNAs accumulated in a polar fashion at the micropylar pole of synergid cells and were also detectable in the micropylar channel that is formed by the integuments of the ovule. Synergid cells are regarded as glandular-like cells that play an active role in secretion. They are the sources of pollen tube attraction signals essential for gametophytic pollen tube guidance (Higashiyama et al., 2001). In dicot plant species such as Torenia and Arabidopsis, defensin-like CRPs known as LUREs are secreted by the synergids as species-specific pollen tube attractants (Okuda et al., 2009; Kanaoka et al., 2011; Takeuchi & Higashiyama, 2012), whereas in maize the secreted EGG APPARATUS1 (ZmEA1) peptide acts as pollen tube attractant (Márton et al., 2005, 2012). Beside pollen tube attraction, synergid cells are also required for pollen tube reception, which involves elaborate communication between the pollen and the receptive synergid cell resulting in pollen tube burst and degeneration of the receptive synergid (Dresselhaus et al., 2016; Higashiyama & Yang 2017). The detection of miR166 and miR390 in Arabidopsis synergid cells and the secretion of these miRNAs into the micropyle suggests hitherto undescribed roles for these accessory cells in secreting small noncoding RNAs, most likely to act in intercellular communication with the arriving pollen tube, or in pathogen defense.

## Author contributions

S.S. conceived the project, J.E. conceived the bioinformatics analyses. M.U., A.B., M.L., C.M., T.H. and S.S. performed experiments and analyzed the data, C.M. generated the small RNA-Seq data. N.S., M.E. and J.E. analyzed the RNA-Seq data. T.D. contributed reagents and analysis tools. S.S. prepared the figures and wrote the manuscript, with input from J.E., M.U. and T.D.

## Acknowledgements

Illumina deep sequencing was carried out at the genomics core facility Center of Excellence for Fluorescent Bioanalytics (KFB, University of Regensburg, Germany). We thank Ingrid Fuchs and Monika Kammerer for their support in tissue culture, Lucija Šoljić for cell isolation and RT-PCR, Nicole Spitzlberger for helping with WISH experiments and Ning Xia with miRNA Northern blots, respectively. This work was supported by the Collaborate Research Centers SFB960, funded by the German Research Foundation (DFG).

## Supplemental Material

### Supplemental Figures

**Figure S1** Pearson’s correlation between samples.

**Figure S2** Expression analysis of gene family members encoding Chromatin-remodeling factors and plant-specific RNA polymerase Pol IV and Pol V subunits.

**Figure S3** Heatmap of egg cell-specific and putative egg cell-specific genes.

**Figure S4** Log2-fold changes of miRNAs which are repressed in the RKD2 egg cell-like callus when compared with the CIM callus.

### Supplemental Tables

**Table S1** List of Oligonucleotides.

**Table S2** Alignment Information for mRNA and small RNA reads from CIM and RKD2-induced callus and for mRNA reads from isolated egg cells.

**Table S3** Differential expression analysis of RNA-seq data generated from RKD2-induced callus and CIM control callus.

**Table S4** Transcripts per million (TPMs) of the two callus types compared with the egg cell

**Table S5** Differential expression of TE sRNA targeted transposable elements and distribution across TE families.

**Table S6** Differentially expressed known miRNAs in the RKD2-induced callus.

**Table S7** Differentially expressed novel miRNAs.

**Table S8** Target prediction and target gene expression of differentially expressed miRNAs in the RKD2-induced callus in comparison to the control callus (CIM).

**Table S9** Differentially in the RKD2-induced callus expressed novel miRNAs with known MIR precursors and their potential target, predicted by psRNAtarget.

**Table S10** Differentially in the RKD2-induced callus expressed novel miRNAs and their potential target, predicted by psRNAtarget.

## References

Allen E, Xie Z, Gustafson AM, Carrington JC. 2005. microRNA-directed phasing during trans-acting siRNA biogenesis in plants. Cell 121: 207–221.

Anders S, Pyl PT, Huber W. 2015. HTSeq - a Python framework to work with high-throughput sequencing data. Bioinformatics 31: 166–169.

Aukermann MJ, Sakai H. 2003. Regulation of flowering time and floral organ identity by a MicroRNA and its APETALA2-like target genes. Plant Cell 15: 2730–2741.

Axtell MJ. 2014. Butter: high-precision genomic alignment of small RNA-seq data. bioRxiv 10.1101/007427.

Baroux C, Raissig MT, Grossniklaus U. (2011). Epigenetic regulation and reprogramming during gamete formation in plants. Current Opinion in Genetics & Development 21: 124–133.

Bolger AM, Lohse M, Usadel B. 2014. Trimmomatic: a flexible trimmer for Illumina sequence data. Bioinformatics 30: 2114–2120.

Borges F, Martienssen RA. 2015. The expanding world of small RNAs in plants. Nat. Rev. Mol. Cell Biol. 16: 727–741.

Borges F, Parent J-S, van Ex F, Wolff P, Martínez G, Köhler C, Martienssen RA. 2018. Transposon-derived small RNAs triggered by miR845 mediate genome dosage response in Arabidopsis. Nature Genetics 50: 186–192.

Brennecke J, Aravin AA, Stark A, Dus M, Kellis M, Sachidanandam R, Hannon GJ. 2007. Discrete small RNA-generating loci as master regulators of transposon activity in Drosophila. Cell 128: 1089–1103.

Clough SJ, Bent AF. 1998. Floral dip: a simplified method for Agrobacterium-mediated transformation of Arabidopsis thaliana. Plant Journal 16: 735–743.

Creasey KM et al. 2014. miRNAs trigger widespread epigenetically activated siRNAs from transposons in *Arabidopsis*. Nature 508: 411–415.

Dai X, Zhao PX. 2011. psRNATarget: a plant small RNA target analysis server. Nucleic Acids Res 39: W155–159.

Dresselhaus T, Sprunck S, Wessel G. 2016. Fertilization mechanisms in flowering plants. Current Biology 26: R125–R139.

Drews GN, Koltunow A. 2011. The Arabidopsis book, the female gametophyte. American Society of Plant Biologists 9: e0155.

Englhart M, Šoljić L, Sprunck S. 2017. Manual isolation of living cells from the Arabidopsis thaliana female gametophyte by micromanipulation. Anja Schmidt (ed.), Plant Germline Development: Methods and Protocols, Methods in Molecular Biology Vol. 1669.

Feng X, Qi Y. 2016. RNAi in Plants: An Argonaute-Centered View. Plant Cell 28: 272–285.

Ghosh DM, Mosiolek M, Bleckmann A, Dresselhaus T, Nodine MD, Maizel A. 2016. Sensitive whole mount in situ localization of small RNAs in plants. Plant Journal 88, 694–702.

Grimson A, Srivastava M, Fahey B, Woodcroft BJ, Chiang HR, King N, Degnan BM, Rokhsar DS, Bartel DP. 2008. Early origins and evolution of microRNAs and Piwi-interacting RNAs in animals. Nature 455, 1193–1197.

Haecker A, Gross-Hardt R, Geiges B, Sarkar A, Breuninger H, Herrmann M, Laux T. 2004. Expression dynamics of WOX genes mark cell fate decisions during early embryonic patterning in Arabidopsis thaliana. Development 131: 657–668.

Hejatko J, Blilou I, Brewer PB, Friml J, Scheres B, Benkova E. 2006. In situ hybridization technique for mRNA detection in whole mount Arabidopsis samples. Nat Protoc 1: 1939–1946.

Higashiyama T, Yabe S, Sasaki N, Nishimura Y, Miyagishima S, Kuroiwa H, Kuroiwa T. 2001. Pollen tube attraction by the synergid cell. Science 293: 1480–1483.

Higashiyama T, Yang WC. 2017. Gametophytic Pollen Tube Guidance: Attractant Peptides, Gametic Controls, and Receptors. Plant Physiol. 173: 112–121.

Ibarra CA, Feng X, Schoft VK, Hsieh TF, Uzawa R, Rodrigues JA, Zemach A, Chumak N, Machlicova A, Nishimura T, Rojas D, Fischer RL, Tamaru H, Zilberman D. 2012. Active DNA demethylation in plant companion cells reinforces transposon methylation in gametes. Science 337, 1360–1364.

Ingouff M, Rademacher S, Foo SH, Šoljić L, Readshaw A, Holec S, Xin N, Lahouze B, Sprunck S, Berger F. (2010). Zygotic resetting of the HISTONE 3 variant repertoire participates in epigenetic reprogramming in Arabidopsis. Curr. Biol. 20: 2137–2143.

Jullien PE, Berger F. (2010). DNA methylation reprogramming during plant sexual reproduction? Trends in Genetics 26: 394–399.

Kanaoka MM, Kawano N, Matsubara Y, Susaki D, Okuda S, Sasaki N, Higashiyama T. 2011. Identification and characterization of TcCRP1, a pollen tube attractant from *Torenia concolor*. Ann Bot. 108: 739–747.

Karimi M, Inze D, Depicker A. 2002. GATEWAY vectors for Agrobacterium-mediated plant transformation. Trends Plant Sci 7: 193–195.

Kersey PJ, Allen JE, Christensen M, Davis P, Falin LJ, Grabmueller C, Hughes DS, Humphrey J, Kerhornou A, Khobova J, et al. 2014. Ensembl Genomes 2013: scaling up access to genome–wide data. Nucleic Acids Res 42 (Database issue): D546–552.

Kim D, Pertea G, Trapnell C, Pimentel H, Kelley R, Salzberg SL. 2013. TopHat2: accurate alignment of transcriptomes in the presence of insertions, deletions and gene fusions. Genome Biol 14: R36.

Koi S, Hisanaga T, Sato K, Shimamura M, Yamato KT, Ishizaki K, Kohchi T, Nakajima K. 2016. An evolutionarily conserved plant RKD factor controls germ cell differentiation. Current Biology 26: 1–7.

Köszegi D, Johnston AJ, Rutten T, Altschmied L, Kumlehn J, Wüst SEJ, Kirioukhova O, Gheyselinck J, Grossniklaus U, Bäumlein H. 2011. Members of the RKD transcription factor family induce an egg cell-like gene expression program. Plant Journal 67: 280–291.

Kozomara A, Griffiths-Jones S. 2014. miRBase: annotating high confidence microRNAs using deep sequencing data. Nucleic Acids Res. 42(Database issue): D68–73.

Lawit SJ, Chamberlin MA, Agee A, Caswell ES, Albertsen MC. 2013. Transgenic manipulation of plant embryo sacs tracked through cell-type specific fluorescent markers: cell labeling, cell ablation, and adventitious embryos. Plant Reproduction 26: 125–137.

Lei J, Sun Y. 2014. miR-PREFeR: an accurate, fast and easy-to-use plant miRNA prediction tool using small RNA-Seq data. Bioinformatics 30:2837–2839.

Liao Y, Smyth GK, Shi W. 2014. featureCounts: an efficient general purpose program for assigning sequence reads to genomic features. Bioinformatics 30: 923–930.

Liu H, Nonomura KI. 2016. A wide reprogramming of histone H3 modifications during male meiosis I in rice is dependent on the Argonaute protein MEL1. J Cell Sci. 129: 3553–3561.

Livak KJ, Schmittgen TD. 2001. Analysis of relative gene expression data using real-time quantitative PCR and the 2(-Delta Delta C(T)) Method. Methods 25: 402–408.

Love MI, Huber W, Anders S. 2014. Moderated estimation of fold change and dispersion for RNA seq data with DESeq2. Genome Biology 15: 550–571.

Márton ML, Cordts S, Broadhvest J, Dresselhaus T. 2005. Micropylar pollen tube guidance by egg apparatus 1 of maize. Science 307: 573–576.

Márton ML, Fastner A, Uebler S, Dresselhaus T. 2012. Overcoming hybridization barriers by the secretion of the maize pollen tube attractant ZmEA1 from Arabidopsis ovules. Curr Biol. 22: 1194– 1198.

Mathieu J, Yant LJ, Murdter F, Kuttner F, Schmid M. 2009. Repression of flowering by the miR172 target SMZ. PLoS Biol 7: e1000148.

Meister G, Landthaler M, Patkaniowska A, Dorsett Y, Teng G, Tuschl T. 2004. Human Argonaute2 mediates RNA cleavage targeted by miRNAs and siRNAs. Mol Cell 15: 185–197.

Nonomura KI. 2018. Small RNA pathways responsible for non-cell-autonomous regulation of plant reproduction. Plant Reprod. 31:21–29.

Nonomura K, Morohoshi A, Nakano M, Eiguchi M, Miyao A, Hirochika H, Kurata N. (2007) A germ cell specific gene of the ARGONAUTE family is essential for the progression of premeiotic mitosis and meiosis during sporogenesis in rice. Plant Cell 19: 2583–2594.

Okuda S, Tsutsui H, Shiina K, Sprunck S, Takeuchi H, Yui R, Kasahara RD, Hamamura Y, Mizukami A, Susaki D, Kawano N, Sakakibara T, Namiki S, Itoh K, Otsuka K, Matsuzaki M, Nozaki H, Kuroiwa T, Nakano A, Kanaoka MM, Dresselhaus T, Sasaki N, Higashiyama T. (2009). Defensin-like polypeptide LUREs are pollen tube attractants secreted from synergid cells. Nature 458: 357 – 361.

Olmedo-Monfil V, Duran-Figueroa N, Arteaga-Vazquez M, Demesa-Arevalo E, Grimanelli D, Slotkin RK, Martienssen RA, Vielle-Calzada JP. 2010. Control of female gamete formation by small RNA pathway in Arabidopsis. Nature 464: 628–632.

Pagnussat GC, Alandete-Saez M, Bowman JL, Sundaresan V. 2009. Auxin-dependent patterning and gamete specification in the Arabidopsis female gametophyte. Science 324: 1684–1689.

Proost S, Mutwil M. 2018. CoNekT: an open-source framework for comparative genomic and transcriptomic network analyses. Nucleic Acids Res 46: W133–W140.

Ramsak Z, Baebler S, Rotter A, Korbar M, Mozetic I, Usadel B, Gruden K. 2014. GoMapMan: integration, consolidation and visualization of plant gene annotations within the MapMan ontology. Nucleic Acids Res 42: D1167–1175.

Resentini F, Cyprys P, Steffen JG, Alter S, Morandini P, Mizzotti C, Lloyd A, Drews GN, Dresselhaus T, Colombo L, Sprunck S, Masiero S. 2017. SUPPRESSOR OF FRIGIDA (SUF4) Supports Gamete Fusion via Regulating Arabidopsis EC1 Gene Expression. Plant Physiol. 173: 155–166.

Rojas-Ríos P, Simonelig M. 2018. piRNAs and PIWI proteins: regulators of gene expression in development and stem cells. Development 145: dev161786. doi: 10.1242/dev.161786.

Rouget C, Papin C, Boureux A, Meunier AC, Franco B, Robine N, Lai EC, Pelisson A, Simonelig M. 2010. Maternal mRNA deadenylation and decay by the piRNA pathway in the early Drosophila embryo. Nature 467: 1128–1132.

Rövekamp M, Bowman JL, Grossniklaus U. 2016. Marchantia MpRKD regulates the gametophyte–sporophyte transition by keeping egg cells quiescent in the absence of fertilization. Current Biology 26: 1–8.

Russell SJ, LaMarre J. 2018. Transposons and the PIWI pathway: genome defense in gametes and embryos. Reproduction 156: R111–R124.

Schmid M, Uhlenhaut NH, Godard F, Demar M, Bressan R, Weigel D, Lohmann JU. 2003. Dissection of floral induction pathways using global expression analysis. Development 130: 6001–6012.

She W, Baroux C. 2015. Chromatin dynamics in pollen mother cells underpin a common scenario at the somatic-to-reproductive fate transition of both the male and female lineages in Arabidopsis. Front. Plant Sci. 6: 294 https://doi.org/10.3389/fpls.2015.00294

Singh M, Goel S, Meeley RB, Dantec C, Parrinello H, Michaud C, Leblanc O, Grimanelli D. 2011. Production of viable gametes without meiosis in maize deficient for an ARGONAUTE protein. Plant Cell 23: 443–458.

Skinner DJ, Sundaresan V. 2018. Recent advances in understanding female gametophyte development. F1000Research 2018, 7 (F1000 Faculty Rev): 804.

Slotkin RK, Vaughn M, Borges F, Tanurdzi M, Becker JD, Feijó JA, Martienssen RA. 2009. Epigenetic reprogramming and small RNA silencing of transposable elements in pollen. Cell 136: 461–472.

Smyth DR, Bowman JL, Meyerowitz EM. 1990. Early flower development in *Arabidopsis*. Plant Cell 2: 755–767.

Sprunck S, Baumann U, Edwards K, Langridge P, Dresselhaus T. 2005. The transcript composition of egg cells changes significantly following fertilization in wheat (*Triticum aestivum* L.). Plant Journal 41: 660–672.

Sprunck S, Rademacher S, Vogler F, Gheyselinck J, Grossniklaus U, Dresselhaus T. 2012. Egg cell-secreted EC1 triggers sperm cell activation during double fertilization. Science 338: 1093–1097.

Takada S, Jürgens G. 2007. Transcriptional regulation of epidermal cell fate in the Arabidopsis embryo. Development 134: 1141–1150

Takeuchi H, Higashiyama T. 2012. A species-specific cluster of defensin-like genes encodes diffusible pollen tube attractants in Arabidopsis. PLoS Biol. 10: e1001449.

Tedeschi F, Rizzo P, Rutten T, Altschmied L, Bäumlein H. 2017. RWP-RK domain-containing transcription factors control cell differentiation during female gametophyte development in Arabidopsis. New Phytologist 213: 1909–1924.

Tucker MR, Okada T, Hu Y, Scholefield A, Taylor JM, Koltunow AMG. (2012) Somatic small RNA pathways promote the mitotic events of megagametogenesis during female reproductive development in Arabidopsis. Development 139: 1399–1404.

Vagin VV, Sigova A, Li C, Seitz H, Gvozdev V, Zamore PD. 2006. A distinct small RNA pathway silences selfish genetic elements in the germline. Science 313, 320–324.

Wang D, Zhang C, Hearn DJ, Kang IH, Punwani JA, Skaggs MI, Drews GN, Schumaker KS, Yadegari R. 2010. Identification of transcription-factor genes expressed in the Arabidopsis female gametophyte. BMC Plant Biol. 10: 110.

Yadegari R, Drews GN. 2004. Female gametophyte development. Plant Cell 16: 133–141.

