## Supplementary material for "Elucidating small RNA pathways in *Arabidopsis thaliana* egg cells": Figure S1

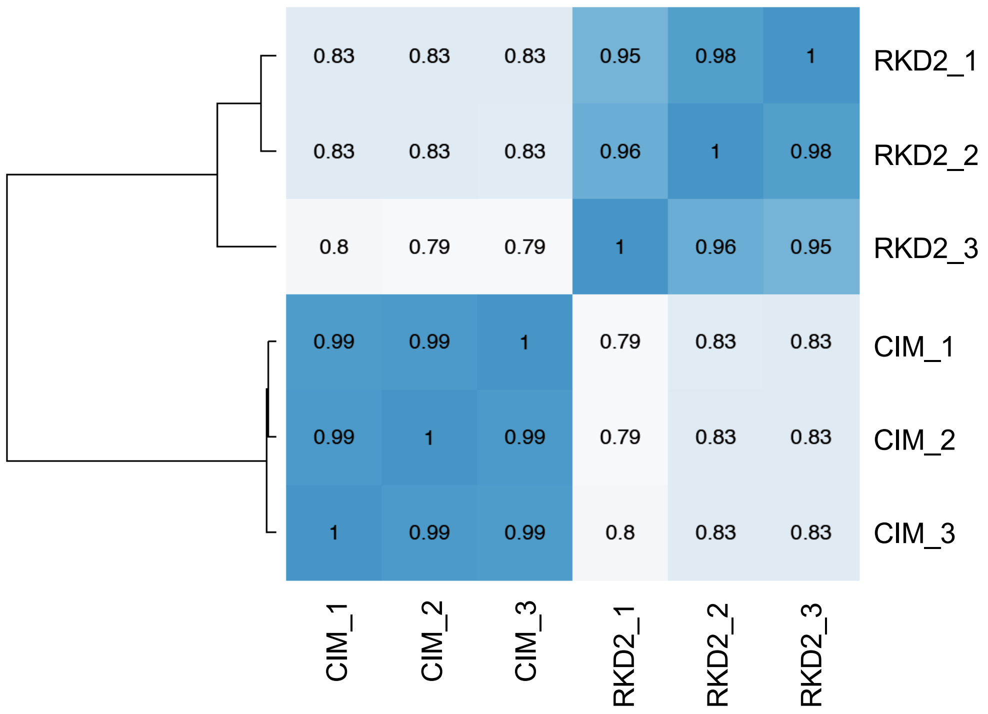


**Figure S1. Pearson's correlation between samples.**

Correlation of samples based on rlog normalized count values of 23,427 genes, which were subsequently tested for differential expression (Love et al., 2014). Pearson correlation values of biological replicates are considerably higher (>0.95) than between the two distinct callus types (0.79 - 0.83), indicating larger differences between the two callus types than between replicates. Numbers (1, 2, 3) indicate biological replicates. Abbreviations: RKD2, RKD2-induced callus; CIM, control callus grown on callus induction medium (CIM) supplemented with auxin and cytokinin.
