## Supplementary material for "Elucidating small RNA pathways in *Arabidopsis thaliana* egg cells": Figure S2

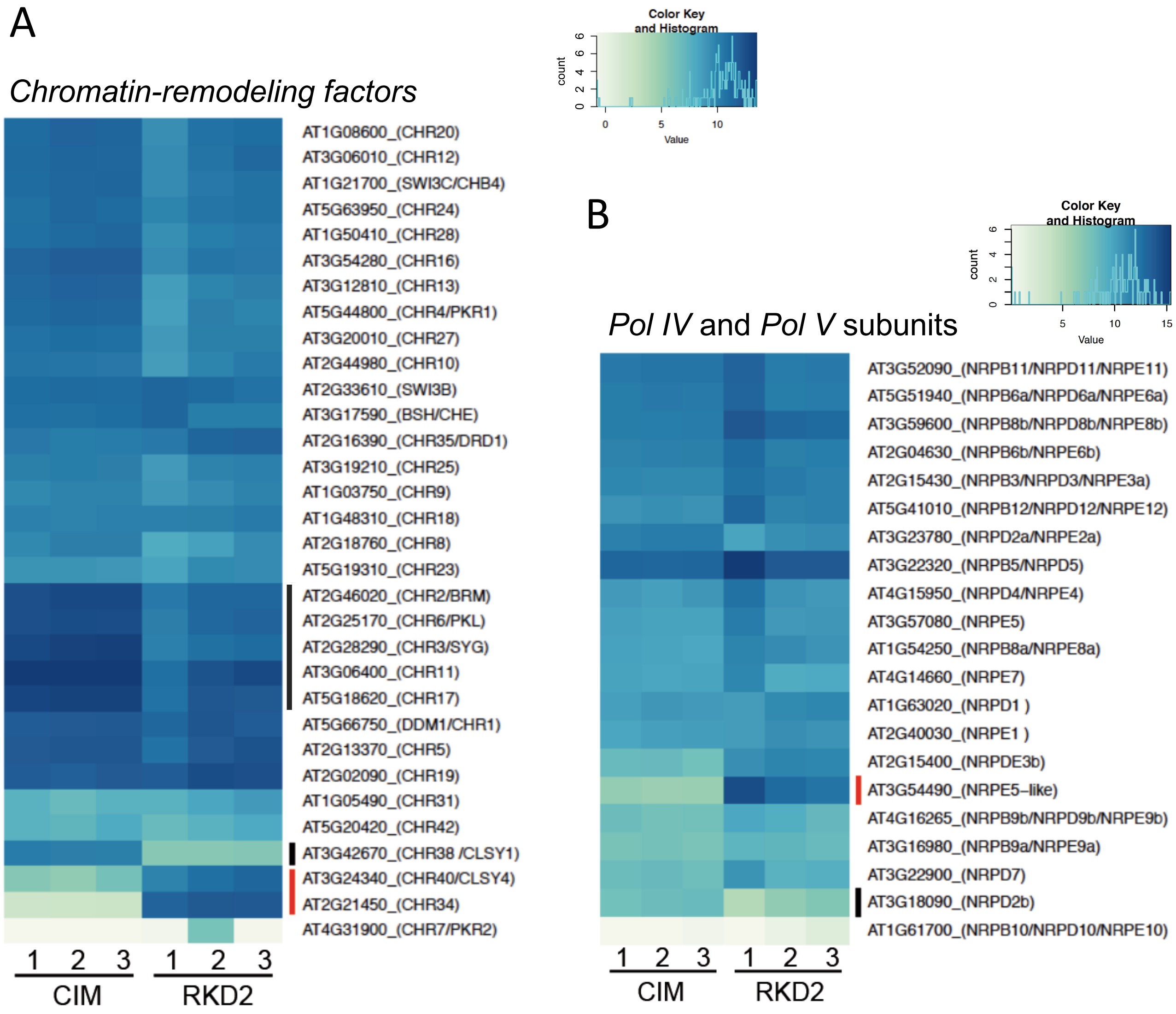


**Figure S2.** Expression analysis of gene family members encoding Chromatin-remodeling factors and plant-specific RNA polymerase Pol IV and Pol V subunits. (a) Heatmap showing expression levels of Arabidopsis SWI/SWF nuclear-localized chromatin remodeling factors (CHRs) in the three biological replicates of the RKD2-induced egg cell-like callus and the hormone-induced control callus (CIM). Note that *CLASSY1* (*CLSY1/CHR38*) is repressed (black bar), while *CLASSY4* (*CHR40)* and *CHR34* are induced in the egg cell-callus (RKD2) (red bar). (b) Heatmap showing expression levels of DNA polymerase IV and V subunits in the three biological replicates of the two callus types. *NRPE5-like* of unknown function, homologous to budding yeast *RPB5*, is strongly induced in the RKD2-induced callus (red bar). Color key indicate normalized and rlog transformed counts.
