## Supplementary material for "Elucidating small RNA pathways in *Arabidopsis thaliana* egg cells": Figure S3

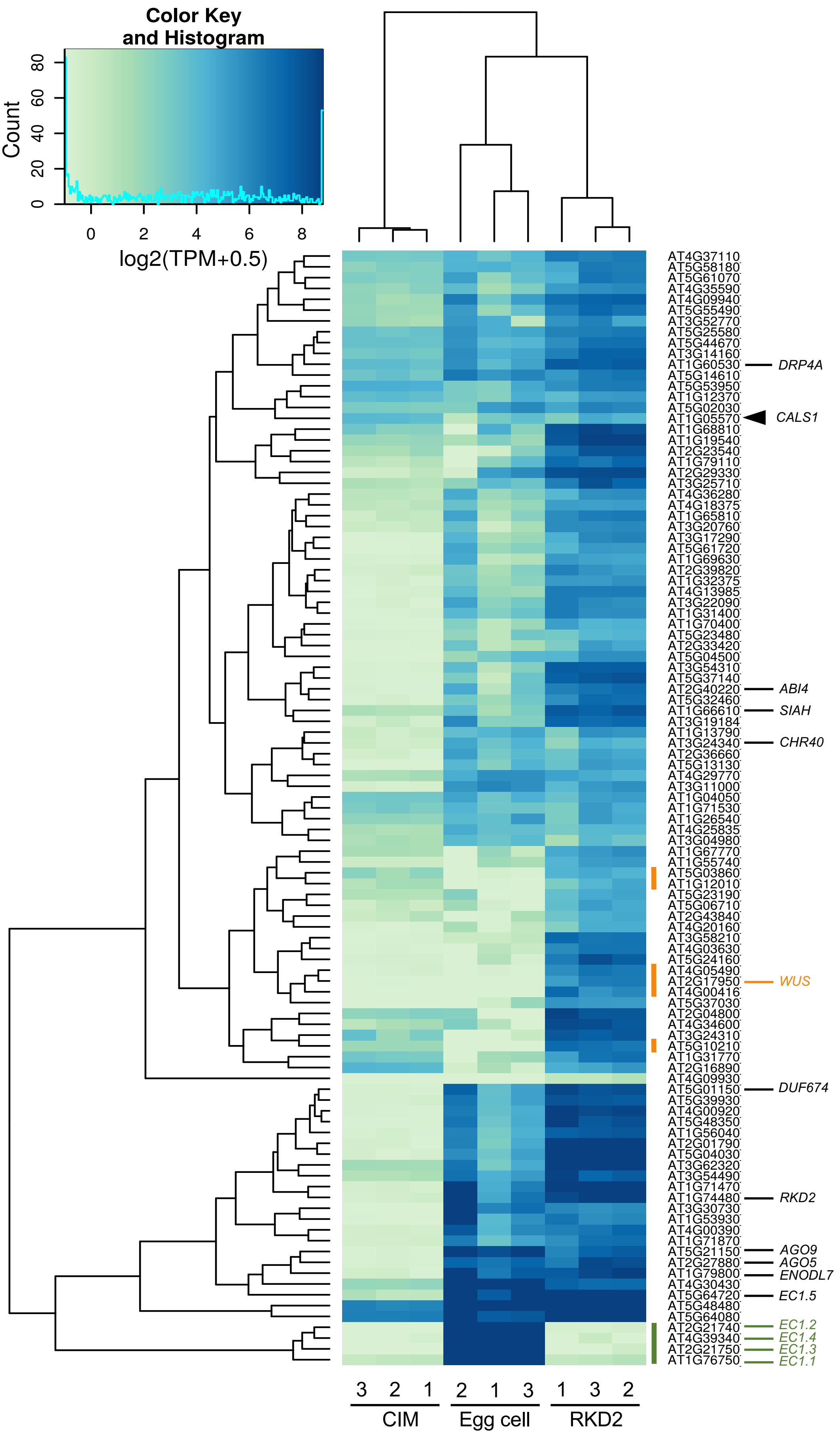

Color Key
and Histogram
Count
0
20
40
60
80
0
2
4
6
8
log2(TPM+0.5)
AT4G37110
AT5G58180
AT5G61070
AT4G35590
AT4G09940
AT5G55490
AT3G52770
AT5G25580
AT5G44670
AT3G14160
AT1G60530
DRP4A
AT5G14610
AT5G53950
AT1G12370
AT5G02030
AT1G05570
CALS1
AT1G68810
AT1G19540
AT2G23540
AT1G79110
AT2G29330
AT3G25710
AT4G36280
AT4G18375
AT1G65810
AT3G20760
AT3G17290
AT5G61720
AT1G69630
AT2G39820
AT1G32375
AT4G13985
AT3G22090
AT1G31400
AT1G70400
AT5G23480
AT2G33420
AT5G04500
AT3G54310
AT5G37140
AT2G40220
ABI4
AT5G32460
AT1G66610
SIAH
AT3G19184
AT1G13790
AT3G24340
CHR40
AT2G36660
AT5G13130
AT4G29770
AT3G11000
AT1G04050
AT1G71530
AT1G26540
AT4G25835
AT3G04980
AT1G67770
AT1G55740
AT5G03860
AT1G12010
AT5G23190
AT5G06710
AT2G43840
AT4G20160
AT3G58210
AT4G03630
AT5G24160
AT4G05490
AT2G17950
WUS
AT4G00416
AT5G37030
AT2G04800
AT4G34600
AT3G24310
AT5G10210
AT1G31770
AT2G16890
AT4G09930
AT5G01150
DUF674
AT5G39930
AT4G00920
AT5G48350
AT1G56040
AT2G01790
AT5G04030
AT3G62320
AT3G54490
AT1G71470
AT1G74480
RKD2
AT3G30730
AT1G53930
AT4G00390
AT1G71870
AT5G21150
AGO9
AT2G27880
AGO5
AT1G79800
ENODL7
AT4G30430
AT5G64720
EC1.5
AT5G48480
AT5G64080
AT2G21740
EC1.2
AT4G39340
EC1.4
AT2G21750
EC1.3
AT1G76750
EC1.1
3 2 1
2 1 3
1 3 2
CIM
Egg cell
RKD2

**Figure S3**. Heatmap of egg cell-specific and putative egg cell-specific genes. The expression profile of 99 genes, previously reported to be upregulated at least seven-fold in the RKD2-induced callus (Kőszegi et al., 2011), was investigated. Four egg cell-specific *EC1* genes (Sprunck et al., 2012) were added to the list. Except one gene (*CALS1*/At1g05570; labeled by black triangle), rlog-normalized counts are considerably higher in the RKD2-induced callus than in the hormone-induced control callus (CIM). Note that *WUSCHEL* (*WUS*) and four other genes (orange bars) are induced in the RKD2-induced callus but not expressed in either of the three egg cell libraries. Except *EC1.5*, no other *EC1* family member is expressed in the RKD2-induced callus (green bar). Statistical analysis was performed using DESeq2 (Love et al., 2014). Color key indicates rlog transformed counts. Expression data of the genes shown in this figure are provided in Table S3.
