## Supplementary material for "Elucidating small RNA pathways in *Arabidopsis thaliana* egg cells": Figure S4

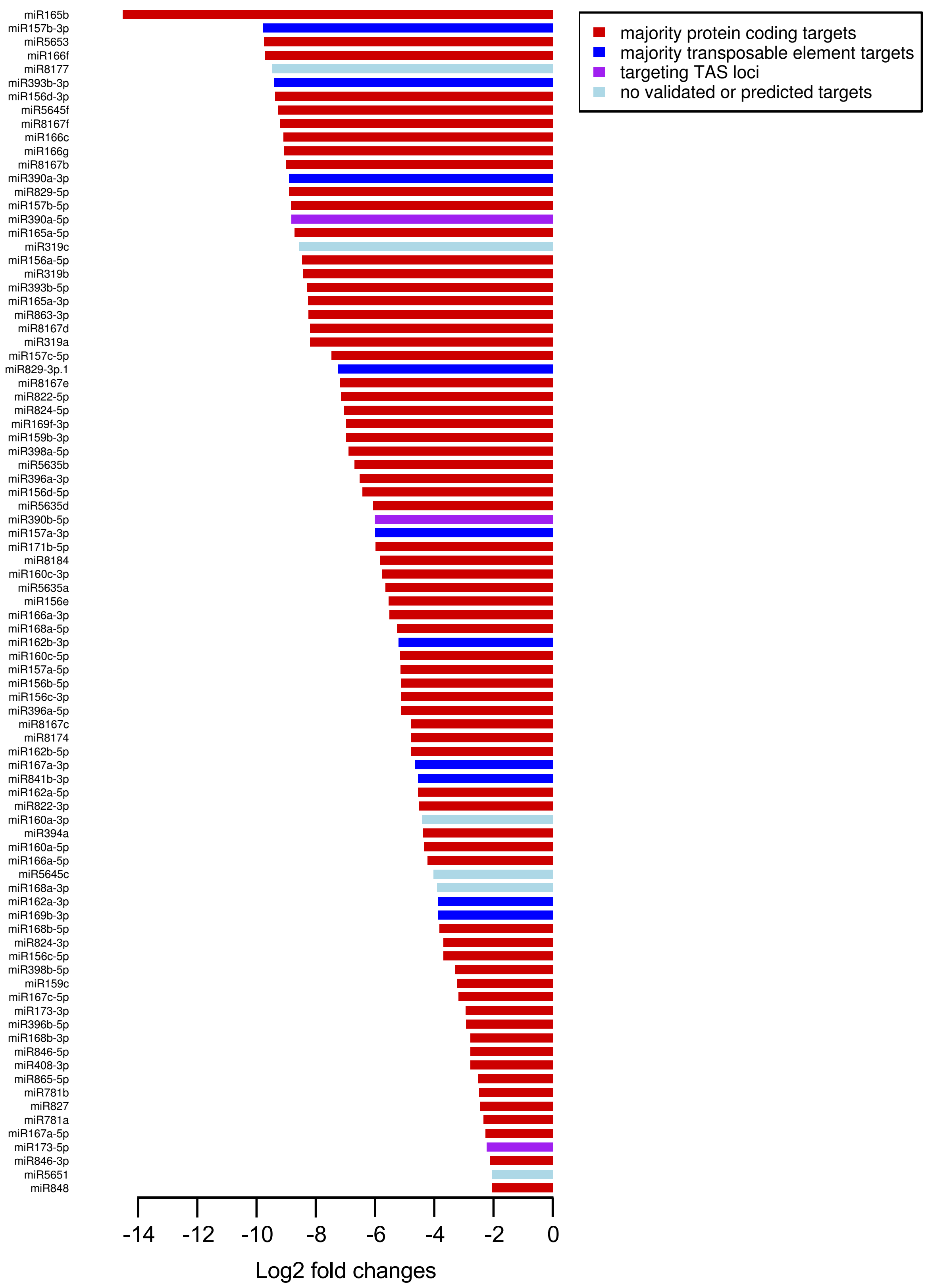


**Figure S4.** Log2-fold changes of miRNAs which are repressed in the RKD2 egg cell-like callus when compared with the CIM callus. Bar colors are based on miRNA targets being mostly protein-coding genes (red), transposable elements (blue), TAS loci (purple), or not having predicted or validated target genes in plant small RNA target analysis (psRNATarget) (light blue).
