## Supplemental Table 1 for "Elucidating small RNA pathways in *Arabidopsis thaliana* egg cells"

**Table S1** List of Oligonucleotides.

| Primer name | Accession | Sequence (5´ to 3´) |
|---|---|---|
| ABI4 RT fw | AT2G40220 | GAGACCGAGTCAAACCGAGTCTAC |
| ABI4 RT rev | | CAGACCGATCCAACATGAACC |
| DRP4A RT fw | AT1G60530 | CATTACATTTATGGTTTTATTCATGCAG |
| DRP4A RT rev | | AGTGTCTGAAACTCCTTCACCAG |
| SINA RT fw | AT1G66610 | CCAAAAGAGGCTACGAGTTCC |
| SINA RT rev | | CATAAGAGAATGTCTCGGTGCAG |
| WOX2 RT fw | AT5G59340 | CGCATGGCTTACTTCAATCG |
| WOX2 RT rev | | TGTTGTGTGACTGTCCGTTTCTC |
| AGO1 for | AT1G48410 | GGCACTTCTCGACCTGCTCAT |
| AGO1 rev | | AGCTGGGAGTGGCCTCACTG |
| AGO2 for | AT1G31280 | ACTTTAAGCAGCCGCGGGGA |
| AGO2 rev | | CTGCGTTATTGGGGCGTTCCGA |
| AGO3 for | AT1G31290 | GGCAATGTGCCTTCAGGTACGGT |
| AGO3 rev | | CGCTCCACGCGACTGCTTAGAATT |
| AGO4 for | AT2G27040 | AAGCACCAGTGCCATTTCTG |
| AGO4 rev | | AACCCCAACAGGCAAACAAG |
| AGO5 for | AT2G27880 | CCCTGAGCAACACGGGAATC |
| AGO5 rev | | TAGTAGCGGGCACGGAATGC |
| AGO6 for | AT2G32940 | GCAGCAGCTCAAGTTGCGCAAT |
| AGO6 for | | AGCTAAACGCAAAAATCGCTGCCG |
| AGO7 for | AT1G69440 | CACGAGCAGGCCAACGCATT |
| AGO7 rev | | AGCCTTCCTCTGTACGCAGCAAG |
| AGO8 for | AT5G21030 | GCGGTTGTGAGCTCCAGAGAG |
| AGO8 rev | | GGGCTTTCGGTTTGGAAGAAC |
| AGO9 for | AT5g21150 | TCGAGGCCCTGATAATGTTCC |
| AGO9 rev | | CTGTGCAGCTGCCAAATGAG |
| AGO10 for | AT5G43810 | ACAGAAGCGTCACCACTCG |
| AGO10 rev | | CGTGCTCGAAATGCTGCAAG |
| AtCB5 fw | AT5G53560 | AGGCGATGAAGTCTTGTTGTCC |
| AtCB5 rev | | CCTTTGGCTTCTTCTAGTCTTTCT |
| qAGO1 for | AT1G48410 | GGCACTTCTCGACCTGCTCAT |
| qAGO1 rev | | AACTGAGCGTGTGCATCTTG |
| qAGO2 for | AT1G31280 | ACTTTAAGCAGCCGCGGGGA |
| qAGO2 rev | | CTGCGTTATTGGGGCGTTCCGA |
| qAGO3 for | AT1G31290 | GGCAATGTGCCTTCAGGTACGGT |
| qAGO3 rev | | CGCTCCACGCGACTGCTTAGAATT |
| qAGO5 for | AT2G27880 | GAACAAGCAGGCCGGCACAT |
| qAGO5 rev | | GCTGGTGGCACAATTGACACAGA |
| qAGO6 for | AT2G32940 | CGTAGCACAACTGCAACTTC |
| qAGO6 for | | GAGGAGGGTTTGGGTTTAGAC |
| qAGO7 for | AT1G69440 | CACGAGCAGGCCAACGCATT |
| qAGO7 rev | | AGCCTTCCTCTGTACGCAGCAAG |
| qAGO9 for | AT5g21150 | TCCAGTCCACACGATAGCT |
| qAGO9 rev | | CCCACAAAAACCGACAGAAT |
| qAGO10 for | AT5G43810 | GTTATACCTATGCGCGGTGCACT |
| qAGO10 rev | | CGTGCTCGAAATGCTGCAAG |

| | | |
|---|---|---|
| qAT5G42470 for | AT5G42470 | ACGCATGGCTGAGCGACTGT |
| qAT5G42470 rev | | TGTTGGAGCAGAGCTTCGTTACA |
| qCHR34 for | AT2G21450 | CAAGCTCGTAGCTGCGGATT |
| qCHR34 rev | | GCATCACCGGAGTGATCAG |
| qCHR40 for | AT3G24340 | CGCATATCCGAACTGGTCTTCT |
| qCHR40 rev | | TTCAGCTTCTCGTGCCGAAC |
| qDCL2 for | AT3G03300 | CGCCTCTCATCATCTCCATAAGC |
| qDCL2 rev | | GCGCCTGCTAGAGATTCTATCAC |
| qEC1.5 for | AT5G64720 | GCGCCGGAAACTTGATGGACT |
| qEC1.5 rev | | GGCGCCGGTGAAGGAGATAAT |
| qNOP56 for | AT1G56110 | CACGCAAGAACGTGGATGTA |
| qNOP56 rev | | TCTTCACCGAGGCATCAACT |
| qNRPE5-L for | AT3G54490 | TGAGGAAGCATGCACTTGAG |
| qNRPE5-L for | | ATCCCCAACAGGCTCTTTAC |
| qSCPL14 for | AT3G12330 | ATGGAGGCCATGGATGATACTCG |
| qSCPL14 rev | | GACCACTTATCCACCTCTTGAAC |
| qSPDS2 for | AT1G70310 | GAAGCCAGTGAGTCTAATCGATAC |
| qSPDS2 for | | GCAAGCAGAAAGCAGCTGAGTG |
| qSWIB/MDM2 for | AT5G23480 | GACCAGCCAATATGGACTGCTTCT |
| qSWIB/MDM2 rev | | CCTTCAGTGATAGAGTCCGTGTTC |
| qUBC21 for | AT5G25760 | TTGTGCCATTGAATTGAACCC |
| qUBC21 rev | | GTCCTGCTTGGACGCTTCAGTCTG |
| qeIF4G for | AT3G60240 | CGGCGATGTTCTTGGGAGTG |
| qeIF4G for | | CCGGTTAGGTGCATGAGGTT |
| **Oligos for cloning** | | |
| AtRKD2 for | AT1G74480 | caccATGGCTGATCACACAACCAAAGAACAGA |
| AtRKD2 rev | | CAAACCACTAGTAAATTCACTTGAGA |
| AGO5p fw | AT2G27880 | caccTCCAAGCCATAGGAAGAGCC |
| AGO5p rev | | TGTAGCGGAAAGCTTCCCAA |
| AGO8p fw | AT5G21030 | caccAGCTTTTAGTATGAAAAAATGCAAGAGATTC |
| AGO8p rev | | TTTCTCGGCCTCCAGATGTTATGC |
| AGO9p fw | AT5g21150 | caccACTTGGGTCATGTAGCCACATCGATGC |
| AGO9p rev | | CTCTGGGATACAAATATCAGAAGGTC |
| DUF674p fw | At5g01150 | caccAGAAACACAAAGTGGAGAGAATCC |
| DUF674p rev | | TAGCGTAATAGATGAAAGAAAGACG |
| ASP_F | At1g31450 | CACCATGGCAACCAAAACTTTTC |
| ASP_R | | TCATAAGTTCCCGGAGCA |
| SBT4_F | At5g59120 | CACCATGGCGACGCTAGCAGCTTCCTCTA |
| SBT4_R | | GTAATCACTAGTATAAACAACAATGGGACTTCTC |
| ENODL7_F | At1g79800 | CCACTGCAACGGCAGCGATCCCATT |
| ENODL7_R | | AATGAAACGAAAGCTAGCAATGGTT |
| AGO1_F | At1g48410 | TCGATTCTACATGGAGCCAGAGACA |
| AGO1_R | | AAGAGTCATAAAGCATCTCATACTCATAGAG |
| **Oligos for Northern blots** | | |
| miR166 | GGGGAATGAAGCCTGGTCCGA | |
| miR172d | CTGCAGCATCATCAAGATTCT | |
| miR390a | GGCGCTATCCCTCCTGAGCTT | |

| | |
|---|---|
| miR781a/b | TAAGTATCCAGAAAACTCTAA |
| miR5653 | GCCAACTCAACTCAACTCAACCCA |
| miR_16 | TTTCCAAAAGTGTAGACAAAG |
| U6 snRNA | TCATCCTTGCGCAGGGGCCA |
| **EXIQON miRCURY LNA enhanced probes, DIG-labeled** | |
| miR390 | 5’-DIG-N-GGCGCTATCCCTCCTGAGCTT-3’-DIG-N |
| miR166 | 5’-N-DIG-GGGGAATGAAGCCTGGTCCGA-3’-DIG-N |
| Ath-miR845a | EXIQON product number 30173-01 |
| Ath-miR172c | EXIQON product number 3006401 |
| Ath-miR95 | custum made: 5'-DIG-N-TGAATCTAATTCCTTCTCTCA |
| Ath-miR8176 | custom made: 5'-DIG-N-TCCCTCTCGCGACCACCGG |
| scramble-miR | EXIQON product number 9904-01 |
