## Supplemental Table 2 for "Elucidating small RNA pathways in *Arabidopsis thaliana* egg cells"

### Table S2. Alignment Information for mRNA and small RNA reads from CIM and RKD2-induced callus and for mRNA reads from isolated egg cells.

**Reference Genome:** ***Arabidopsis thaliana*****, TAIR 10** (ftp://ftp.arabidopsis.org/home/tair/Sequences/whole_chromosomes/TAIR10_chr*.fas; 22.02.2011)

#### A. mRNA analysis

Software: Tophat2

| | Left | | | Right | | | | Aligned pairs | | | |
|---|---|---|---|---|---|---|---|---|---|---|---|
| library | Input | Mapped unique | Mapped multiple | Input | Mapped unique | Mapped multiple | Overall read mapping rate | Mapped unique | Mapped multiple | Mapped discorda | Concordant pair alignment rate |
| CIM-1 | 59575423 | 49555346 | 6150541 | 59575423 | 49333640 | 6127077 | 93.3% | 47950394 | 5975325 | 903238 | 89.0% |
| CIM-2 | 63382444 | 55105305 | 5718724 | 63382444 | 54854097 | 5697143 | 95.7% | 53343033 | 5553662 | 790420 | 91.7% |
| CIM-3 | 59457993 | 51724265 | 5576229 | 59457993 | 51422237 | 5551008 | 96.1% | 50050724 | 5424931 | 1112154 | 91.4% |
| RKD2-1 | 53697155 | 47792273 | 3456882 | 53697155 | 47526886 | 3440690 | 95.2% | 46264002 | 3364501 | 543989 | 91.4% |
| RKD2-2 | 54284662 | 48361468 | 3933941 | 54284662 | 48086384 | 3914862 | 96.1% | 46809852 | 3828982 | 911409 | 91.6% |
| RKD2-3 | 52444623 | 47103697 | 3256023 | 52444623 | 46808215 | 3238981 | 95.7% | 45479374 | 3159461 | 856378 | 91.1% |
| Egg_Pool_1 | 47409104 | 40663329 | 1506939 | 47409104 | 40669810 | 1507271 | 89.0% | 38316006 | 1400487 | 111391 | 83.5% |
| Egg_Pool_2 | 56700468 | 22541234 | 1008601 | 56700468 | 22559422 | 1005387 | 41.5% | 20956321 | 860643 | 57111 | 38.4% |
| Egg_Pool_3 | 50644012 | 25479370 | 1154971 | 50644012 | 25470536 | 1151880 | 52.6% | 24122110 | 1045937 | 63111 | 49.6% |

#### B. small RNA analysis

Software: Trimmomatic

| library | input_reads | surviving | surviving(%) | dropped | dropped(%) |
|---|---|---|---|---|---|
| CIM-1 | 32291776 | 28968013 | (89.71%) | 3323763 | (10.29%) |
| CIM-2 | 31847955 | 29221993 | (91.75%) | 2625962 | (8.25%) |
| CIM-3 | 28208451 | 26224301 | (92.97%) | 1984150 | (7.03%) |
| RKD2-1 | 25765018 | 20198410 | (78.39%) | 5566608 | (21.61%) |
| RKD2-2 | 24888241 | 16981594 | (68.23%) | 7906647 | (31.77%) |
| RKD2-3 | 26581195 | 19074366 | (71.76%) | 7506829 | (28.24%) |

Software: Butter 0.3.3

| **library** | **readlength** | **Input** | **Unmapped** | **Uniquely mapped reads** | **Multi mapped reads** | **Multi mapped randomly assigned** | **Multi mapped density assigned** |
|---|---|---|---|---|---|---|---|
| CIM-1 | 18-24 | 13.659.320 | (14.2%) | (48.6%) | (37.2%) | (1.6%) | (35.6%) |
| CIM-1 | 25-30 | 1.220.945 | (29.0%) | (27.4%) | (43.5%) | (11.0%) | (32.5%) |
| CIM-1 | 31-50 | 14.087.748 | (56.9%) | (36.5%) | (6.6%) | (0.5%) | (6.1%) |
| | **total** | **28.968.013** | **(35.58%)** | **(41.84%)** | **(22.58%)** | **(1.44%)** | **(21.14%)** |
| CIM-2 | 18-24 | 12.084.682 | (14.0%) | (48.2%) | (37.8%) | (1.4%) | (36.4%) |
| CIM-2 | 25-30 | 1.218.315 | (29.1%) | (30.9%) | (40.0%) | (10.9%) | (29.0%) |
| CIM-2 | 31-50 | 15.918.996 | (60.2%) | (33.3%) | (6.5%) | (0.5%) | (6.0%) |
| | **total** | **29.221.993** | **(39.81%)** | **(39.37%)** | **(20.82%)** | **(1.32%)** | **(19.5%)** |
| CIM-3 | 18-24 | 10.477.961 | (14.3%) | (49.6%) | (36.2%) | (1.3%) | (34.9%) |
| CIM-3 | 25-30 | 891.556 | (30.8%) | (31.2%) | (38.0%) | (9.8%) | (28.3%) |
| CIM-3 | 31-50 | 14.854.784 | (49.7%) | (42.6%) | (7.7%) | (0.5%) | (7.3%) |
| | **total** | **26.224.301** | **(34.88%)** | **(44.99%)** | **(20.13%)** | **(1.1%)** | **(19.03%)** |
| RKD2-1 | 18-24 | 3.617.686 | (8.3%) | (20.7%) | (71.0%) | (12.9%) | (58.1%) |
| RKD2-1 | 25-30 | 2.901.304 | (12.4%) | (9.9%) | (77.7%) | (15.5%) | (62.2%) |
| RKD2-1 | 31-50 | 13.679.420 | (80.8%) | (9.5%) | (9.7%) | (1.5%) | (8.1%) |
| | **total** | **20.198.410** | **(58.02%)** | **(11.56%)** | **(30.42%)** | **(5.59%)** | **(24.83%)** |
| RKD2-2 | 18-24 | 5.321.437 | (8.8%) | (21.3%) | (69.9%) | (12.8%) | (57.1%) |
| RKD2-2 | 25-30 | 3.072.394 | (12.5%) | (10.4%) | (77.1%) | (16.5%) | (60.6%) |
| RKD2-2 | 31-50 | 8.587.763 | (91.7%) | (4.0%) | (4.3%) | (0.8%) | (3.4%) |
| | **total** | **16.981.594** | **(51.41%)** | **(10.59%)** | **(38%)** | **(7.41%)** | **(30.59%)** |
| RKD2-3 | 18-24 | 5.460.894 | (7.3%) | (17.4%) | (75.3%) | (14.9%) | (60.4%) |
| RKD2-3 | 25-30 | 3.711.206 | (13.0%) | (10.1%) | (76.9%) | (16.3%) | (60.7%) |
| RKD2-3 | 31-50 | 9.902.266 | (94.7%) | (2.0%) | (3.3%) | (0.7%) | (2.7%) |
| | **total** | **19.074.366** | **(53.78%)** | **(7.97%)** | **(38.26%)** | **(7.78%)** | **(30.48%)** |
