## Supplemental Table 6 for "Elucidating small RNA pathways in *Arabidopsis thaliana* egg cells"

**Table S6.** Differentially expressed known miRNAs in the RKD2-induced callus. Log2-fold changes and miRNA counts in the three replicates from the control callus (CIM) and the RKD2-induced callus are shown.

| | log2FC.miRNA | counts.miRNA.CIM_1 | counts.miRNA.CIM_2 | counts.miRNA.CIM_3 | counts.miRNA.RKD2_1 | counts.miRNA.RKD2_2 | counts.miRNA.RKD2_3 |
|---|---|---|---|---|---|---|---|
| miR172d-3p | 9,2 | 1,36 | 0 | 1,85 | 920,35 | 542,65 | 687,92 |
| miR831-3p | 6,1 | 0 | 0,83 | 0 | 17,9 | 26,52 | 31,5 |
| miR866-5p | 6,1 | 31,94 | 34,08 | 42,53 | 2215,11 | 2314,56 | 2905,38 |
| miR845a | 5,3 | 4,76 | 1,66 | 2,77 | 38,78 | 129,74 | 219,23 |
| miR172d-5p | 4,7 | 2,04 | 0 | 0 | 31,32 | 26,52 | 11,34 |
| miR158a-5p | 4,5 | 246 | 364,93 | 273,69 | 5889,05 | 6688,93 | 6997,6 |
| miR866-3p | 4,2 | 42,13 | 25,77 | 36,06 | 593,68 | 566,33 | 696,74 |
| miR5640 | 3,7 | 138,63 | 136,33 | 107,26 | 1572,2 | 1645,01 | 1629,08 |
| miR8176 | 3 | 14,27 | 11,98 | 11,1 | 99,94 | 65,35 | 136,07 |
| miR848 | -2,1 | 55,04 | 73,15 | 57,33 | 8,95 | 13,26 | 21,42 |
| miR5651 | -2,1 | 515,79 | 509,57 | 519,64 | 120,82 | 138,27 | 105,83 |
| miR846-3p | -2,1 | 6185,39 | 8915,47 | 7756,65 | 1351,44 | 1940,48 | 1925,16 |
| miR173-5p | -2,2 | 208,63 | 271,83 | 194,17 | 49,22 | 55,88 | 36,54 |
| miR167a-5p | -2,3 | 112,13 | 194,52 | 176,6 | 29,83 | 34,09 | 34,02 |
| miR781a | -2,3 | 87,66 | 93,93 | 71,2 | 19,39 | 15,15 | 15,12 |
| miR827 | -2,5 | 687,04 | 1376,6 | 1405,42 | 244,63 | 276,54 | 97,01 |
| miR781b | -2,5 | 89,7 | 83,96 | 92,46 | 16,41 | 13,26 | 17,64 |
| miR865-5p | -2,5 | 137,27 | 150,46 | 146,09 | 26,85 | 28,41 | 18,9 |
| miR408-3p | -2,8 | 7531,6 | 13862,41 | 12252,16 | 1585,63 | 1929,12 | 1333 |
| miR846-5p | -2,8 | 1004,39 | 1130,54 | 1116,02 | 125,3 | 180,88 | 158,75 |
| miR168b-3p | -2,8 | 219,5 | 317,55 | 289,41 | 55,19 | 38,83 | 25,2 |
| miR396b-5p | -2,9 | 3531,69 | 4475,61 | 3595,85 | 557,88 | 714,07 | 220,49 |
| miR173-3p | -2,9 | 1628,23 | 1872,04 | 1464,6 | 234,19 | 222,55 | 185,21 |
| miR167c-5p | -3,2 | 90,38 | 147,97 | 123,9 | 13,42 | 15,15 | 10,08 |
| miR159c | -3,2 | 32,62 | 42,4 | 32,36 | 1,49 | 0,95 | 8,82 |
| miR398b-5p | -3,3 | 642,19 | 753,97 | 632,44 | 99,94 | 51,14 | 54,18 |
| miR156c-5p | -3,7 | 4152,81 | 5558,76 | 4783,07 | 736,88 | 214,03 | 97,01 |
| miR824-3p | -3,7 | 960,9 | 1448,09 | 1347,17 | 80,55 | 73,87 | 132,29 |
| miR168b-5p | -3,8 | 1370 | 1794,73 | 1780,82 | 153,64 | 117,43 | 73,08 |
| miR169b-3p | -3,9 | 15,63 | 27,43 | 36,06 | 2,98 | 0 | 2,52 |
| miR162a-3p | -3,9 | 683,64 | 694,95 | 412,38 | 35,8 | 62,5 | 18,9 |
| miR168a-3p | -3,9 | 847,42 | 1086,48 | 800,72 | 70,11 | 53,98 | 56,7 |
| miR5645c | -4,0 | 30,58 | 44,06 | 49,93 | 4,47 | 2,84 | 0 |
| miR166a-5p | -4,2 | 655,78 | 760,62 | 644,46 | 41,77 | 44,51 | 21,42 |
| miR160a-5p | -4,3 | 94,46 | 189,53 | 141,47 | 11,93 | 8,52 | 0 |
| miR394a | -4,4 | 22,43 | 36,58 | 103,56 | 2,98 | 1,89 | 2,52 |
| miR160a-3p | -4,4 | 20,39 | 39,9 | 25,89 | 2,98 | 0,95 | 0 |
| miR822-3p | -4,5 | 116,21 | 190,36 | 138,69 | 2,98 | 7,58 | 7,56 |
| miR162a-5p | -4,5 | 48,93 | 65,67 | 36,98 | 2,98 | 0,95 | 2,52 |
| miR841b-3p | -4,6 | 39,41 | 39,07 | 12,02 | 0 | 2,84 | 0 |
| miR167a-3p | -4,6 | 428,13 | 1464,71 | 969,93 | 32,82 | 44,51 | 34,02 |
| miR162b-5p | -4,8 | 16,99 | 39,9 | 21,27 | 0 | 0 | 2,52 |
| miR8174 | -4,8 | 149,5 | 133,84 | 159,03 | 7,46 | 5,68 | 2,52 |
| miR8167c | -4,8 | 35,34 | 20,78 | 22,19 | 1,49 | 0 | 1,26 |
| miR396a-5p | -5,1 | 27731,63 | 30502,11 | 27249,52 | 636,94 | 709,33 | 1099,91 |
| miR156c-3p | -5,1 | 880,71 | 990,05 | 782,23 | 43,26 | 20,83 | 12,6 |
| miR156b-5p | -5,1 | 6554,39 | 5270,31 | 5104,84 | 114,86 | 219,71 | 138,59 |
| miR157a-5p | -5,1 | 579,67 | 941,01 | 1484,02 | 14,92 | 11,36 | 52,92 |
| miR160c-5p | -5,1 | 254,16 | 389,04 | 400,36 | 5,97 | 7,58 | 15,12 |
| miR162b-3p | -5,2 | 1790,65 | 2058,25 | 1939,86 | 26,85 | 29,36 | 97,01 |
| miR168a-5p | -5,3 | 12686,77 | 14258,09 | 13459,72 | 359,49 | 371,24 | 313,72 |
| miR166a-3p | -5,5 | 221636,24 | 329912,12 | 288884,15 | 5444,54 | 8242,07 | 4346,73 |
| miR156e | -5,5 | 553,84 | 480,48 | 634,29 | 20,88 | 9,47 | 0 |
| miR5635a | -5,6 | 29,22 | 24,11 | 25,89 | 0 | 0,95 | 0 |
| miR160c-3p | -5,8 | 90,38 | 108,07 | 88,76 | 2,98 | 0,95 | 1,26 |
| miR8184 | -5,8 | 166,49 | 172,07 | 169,21 | 5,97 | 0,95 | 2,52 |
| miR171b-5p | -6,0 | 78,15 | 104,74 | 69,35 | 0 | 1,89 | 1,26 |

| | | | | | | | |
|---|---|---|---|---|---|---|---|
| miR157a-3p | -6,0 | 173,29 | 239,41 | 399,44 | 2,98 | 1,89 | 7,56 |
| miR390b-5p | -6,0 | 67,96 | 42,4 | 155,34 | 0 | 0,95 | 2,52 |
| miR5635d | -6,1 | 35,34 | 37,41 | 34,21 | 0 | 0,95 | 0 |
| miR156d-5p | -6,4 | 6716,81 | 10633,72 | 7817,68 | 43,26 | 85,23 | 156,23 |
| miR396a-3p | -6,5 | 1321,75 | 1474,69 | 1168,72 | 17,9 | 8,52 | 17,64 |
| miR5635b | -6,7 | 147,47 | 126,35 | 147,94 | 1,49 | 0 | 2,52 |
| miR398a-5p | -6,9 | 198,43 | 236,08 | 193,25 | 0 | 0,95 | 3,78 |
| miR159b-3p | -7,0 | 8035,16 | 11024,42 | 12240,14 | 83,53 | 69,13 | 94,49 |
| miR169f-3p | -7,0 | 59,12 | 87,28 | 61,95 | 0 | 0,95 | 0 |
| miR824-5p | -7,0 | 375,12 | 564,44 | 567,72 | 2,98 | 0,95 | 7,56 |
| miR822-5p | -7,2 | 1149,82 | 1276,84 | 1245,47 | 5,97 | 6,63 | 12,6 |
| miR8167e | -7,2 | 73,39 | 102,25 | 68,42 | 0 | 0 | 1,26 |
| miR829-3p.1 | -7,3 | 507,63 | 743,16 | 697,16 | 7,46 | 1,89 | 3,78 |
| miR157c-5p | -7,5 | 7880,22 | 9006,91 | 7985,03 | 28,34 | 50,19 | 57,96 |
| miR319a | -8,2 | 2506,23 | 3280,23 | 3644,86 | 4,47 | 6,63 | 20,16 |
| miR8167d | -8,2 | 50,97 | 43,23 | 56,4 | 0 | 0 | 0 |
| miR863-3p | -8,2 | 40,09 | 72,32 | 46,23 | 0 | 0 | 0 |
| miR165a-3p | -8,3 | 53387,87 | 75399,48 | 60884,48 | 150,66 | 107,96 | 340,18 |
| miR393b-5p | -8,3 | 58,44 | 45,72 | 58,25 | 0 | 0 | 0 |
| miR319b | -8,4 | 32137,24 | 36915,43 | 40529,86 | 64,14 | 35,99 | 187,73 |
| miR156a-5p | -8,5 | 61,84 | 61,51 | 61,95 | 0 | 0 | 0 |
| miR319c | -8,6 | 10140,45 | 10423,41 | 11182,37 | 17,9 | 8,52 | 49,14 |
| miR165a-5p | -8,7 | 65,92 | 83,13 | 78,59 | 0 | 0 | 0 |
| miR390a-5p | -8,8 | 1603,77 | 1872,87 | 1686,51 | 1,49 | 1,89 | 7,56 |
| miR157b-5p | -8,8 | 10896,12 | 10785,01 | 7935,1 | 20,88 | 4,74 | 37,8 |
| miR829-5p | -8,9 | 432,2 | 482,97 | 508,54 | 1,49 | 0 | 1,26 |
| miR390a-3p | -8,9 | 82,23 | 93,93 | 89,69 | 0 | 0 | 0 |
| miR8167b | -9,0 | 92,42 | 70,66 | 126,67 | 0 | 0 | 0 |
| miR166g | -9,1 | 1042,45 | 319,21 | 462,31 | 0 | 0 | 2,52 |
| miR166c | -9,1 | 2891,54 | 6442,41 | 3236,18 | 2,98 | 13,26 | 3,78 |
| miR8167f | -9,2 | 112,13 | 96,43 | 124,82 | 0 | 0 | 0 |
| miR5645f | -9,3 | 116,21 | 113,89 | 127,6 | 0 | 0 | 0 |
| miR156d-3p | -9,4 | 127,76 | 149,63 | 108,18 | 0 | 0 | 0 |
| miR393b-3p | -9,4 | 125,72 | 126,35 | 139,62 | 0 | 0 | 0 |
| miR8177 | -9,5 | 154,26 | 137,16 | 121,13 | 0 | 0 | 0 |
| miR166f | -9,7 | 10664,39 | 20761,19 | 14658,95 | 19,39 | 25,57 | 6,3 |
| miR5653 | -9,7 | 785,58 | 922,72 | 885,79 | 0 | 0 | 2,52 |
| miR157b-3p | -9,8 | 3222,49 | 2708,31 | 2063,76 | 2,98 | 0,95 | 5,04 |
| miR165b | -14,52 | 5588,05 | 9671,93 | 6034,08 | 0 | 0 | 0 |
